# Vacuolar H^+^-ATPase Determines Daughter Cell Fates through Asymmetric Segregation of the Nucleosome Remodeling and Deacetylase Complex

**DOI:** 10.1101/2023.06.25.546476

**Authors:** Zhongyun Xie, Yongping Chai, Zhiwen Zhu, Zijie Shen, Zhengyang Guo, Zhiguang Zhao, Long Xiao, Zhuo Du, Guangshuo Ou, Wei Li

## Abstract

Asymmetric cell divisions (ACDs) generate two daughter cells with identical genetic information but distinct cell fates through epigenetic mechanisms. However, the process of partitioning different epigenetic information into daughter cells remains unclear. Here, we demonstrate that the nucleosome remodeling and deacetylase (NuRD) complex is asymmetrically segregated into the surviving daughter cell rather than the apoptotic one during ACDs in *Caenorhabditis elegans*. The absence of NuRD triggers apoptosis via the EGL-1-CED-9-CED-4-CED-3 pathway, while an ectopic gain of NuRD enables apoptotic daughter cells to survive. We identify the vacuolar H^+^–adenosine triphosphatase (V-ATPase) complex as a crucial regulator of NuRD’s asymmetric segregation. V-ATPase interacts with NuRD and is asymmetrically segregated into the surviving daughter cell. Inhibition of V-ATPase disrupts cytosolic pH asymmetry and NuRD asymmetry. We suggest that asymmetric segregation of V-ATPase may cause distinct acidification levels in the two daughter cells, enabling asymmetric epigenetic inheritance that specifies their respective life-versus-death fates.

## Introduction

Asymmetric cell division (ACD) gives rise to two daughter cells that possess identical genetic material but distinct cell fates, playing a crucial role in both development and tissue homeostasis (Royall and Jessberger, 2021; Sunchu and Cabernard, 2020; Venkei and Yamashita, 2018; Wooten et al., 2020; Zion et al., 2020). Both extrinsic and intrinsic mechanisms determine distinct daughter cell fates after ACD. While extrinsic mechanisms, such as exposure to signaling cues from the local niche, have been extensively studied and are known to define stem cell fate (Morrison and Spradling, 2008), the intrinsic mechanisms are more complex and largely unresolved, despite several proteins, RNA molecules, and organelles having been implicated in the regulation of some types of ACD (Sunchu and Cabernard, 2020; Zion et al., 2020). Epigenetic mechanisms play a crucial role in guiding the two daughter cells towards establishing differential gene expression profiles, ultimately defining their unique cell identities (Allis and Jenuwein, 2016; Escobar et al., 2021; Stewart-Morgan et al., 2020). During the ACD of *Drosophila* male germline stem cells (GSCs), pre-existing and newly synthesized histones H3 and H4 are asymmetrically segregated towards the stem daughter cell and the differentiating daughter cell, respectively (Wooten et al., 2020). Nevertheless, the extent to which epigenetic information is asymmetrically inherited through ACD in other organisms and the mechanism by which this process occurs remains elusive.

*Caenorhabditis elegans* represents a valuable model for investigating ACD, given its invariant cell lineage and conserved mechanisms of ACD. During hermaphrodite development in *C. elegans*, 131 somatic cells undergo programmed cell death, with the majority produced through ACD that create a large cell programmed for survival and a small cell programmed to die (Sulston and Horvitz, 1977; Sulston et al., 1983a) The opposing cell fates of daughter cells, i.e., to live or die, offer a compelling experimental system for investigating how epigenetic inheritance determines life versus death decisions during ACD. It is noteworthy that 105 of the 131 apoptotic cells arise from neuronal lineages, of which the Q neuroblast represents a tractable system for studying ACD and apoptosis at single-cell resolution (Ellis and Horvitz, 1986; Hedgecock et al., 1983; Ou et al., 2010; Sulston and Horvitz, 1977). During the first larval stage, Q neuroblasts on the left (QL) and right (QR) undergo three rounds of asymmetric divisions, which produce three different neurons and two apoptotic cells (Q.aa and Q.pp), respectively (Figure 1A).

**Figure 1:**
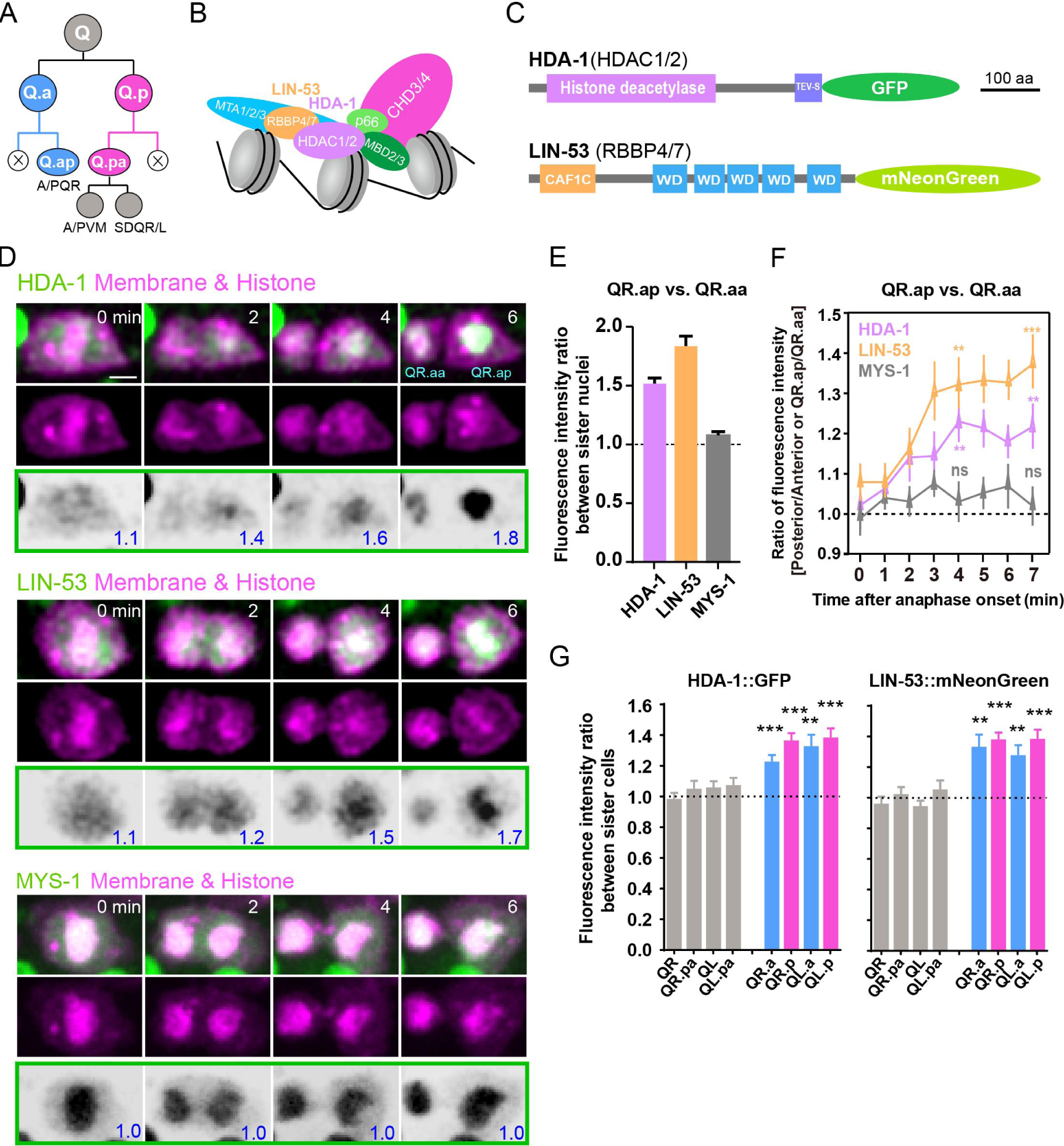
Asymmetric segregation of NuRD during ACDs of *C. elegans* Q neuroblast. (A) Schematic of the Q neuroblast lineages. QL or QR neuroblast each generates three neurons and two apoptotic cells (Q.aa/Q.pp, X). QL produces PQR, PVM, and SDQL. QR produces AQR, AVM, and SDQR. (B) A model of the NuRD complex composition (Bracken et al., 2019; Lai and Wade, 2011). (C) Protein domain structure of the GFP-tagged HDAC1/2 (HDA-1) or mNeonGreen-tagged RBBP4/7 (LIN-53). CAF1C: histone-binding protein RBBP4 or subunit C of CAF1 complex; WD: WD40 repeat tryptophan-aspartate domain. Scale bar: 100 amino acids. (D) Representative images of endogenous HDA-1::GFP, LIN-53::mNeonGreen and overexpressed MYS-1::GFP during ACDs of QR.a. In each panel, the top row shows merged images, the middle row shows mCherry-tagged plasma membrane and histone, and the bottom row shows inverted fluorescence images of GFP/mNeonGreen. The anterior of the cell is on the left. The GFP/mNeonGreen fluorescence intensity ratios between posterior and anterior chromatids, and between QR.ap and QR.aa nuclei, are shown in blue at the lower right corner of inverted fluorescence images. Other frames are in Figure S3A, and the full movies are in Supplementary Movie S3-S5. Scale bar: 2 µm. (E) Quantification of HDA-1::GFP, LIN-53::mNeonGreen and MYS-1::GFP fluorescence intensity ratio between QR.ap and QR.aa nuclei. Data are presented as mean ± SEM. N = 10–12. (F) Quantification of HDA-1::GFP (magenta), LIN-53::mNeonGreen (orange) and MYS-1::GFP (gray) fluorescence intensity ratios between the posterior and anterior half of QR.a or between QR.ap and QR.aa. Anaphase onset is defined as the last frame without chromatids segregation. Data are presented as mean ± SEM. N = 10–12. Statistical significance is determined by a one-sample t-test with 1 as the theoretical mean. ** p < 0.01, *** p < 0.001, ns: not significant. (G) Quantification of HDA-1::GFP (left) and LIN-53::mNeonGreen (right) fluorescence intensity ratios between the large and small daughters of cells shown on the X-axis. Data are presented as mean ± SEM. N = 7–14. Statistical significance is determined by a one-sample t-test with 1 as the theoretical mean. ** p < 0.01, *** p < 0.001.

In the nematode, the classic apoptotic pathway initiates upon activation of the BH3-only protein EGL-1 in the cells that are fated to die. EGL-1 binds to the anti-apoptotic Bcl-2-like protein CED-9, which facilitates the release of the pro-apoptotic protein CED-4, leading to caspase CED-3 activation. Activated caspase promotes the exposure of phosphatidylserine (PS) on the surface of apoptotic cells, which triggers the phagocytosis of apoptotic cells via partially redundant signaling pathways such as the CED-1/CED-6/CED-7 pathway and the CED-2/CED-5/CED-12 pathway. Proper transcriptional regulation of the *egl-1* gene is vital for life-versus-death decisions in *C. elegans* (Conradt and Horvitz, 1998; Nehme and Conradt, 2008). Although the cell-specific transcriptional regulator has been identified in certain cells, including the sexually dimorphic HSN and CEM neurons, M4 motor neuron, NSM neuron, P11.aaap cell, ABpl/rpppapp, and the tail spike cell (Hirose et al., 2010; Jiang and Wu, 2014; Nehme and Conradt, 2008), the general upstream regulators of *egl-1* at the chromatin remodeling or epigenetic level are unclear. Two NSM neuroblasts undergo ACDs to each generate a larger NSM cell programed to survive and a smaller NSM sister cell programed to die (Sulston et al., 1983b). The PIG-1 kinase-dependent asymmetric partitioning of the Snail-like transcription factor, CES-1, represses *egl-1* transcription in the larger NSM sister cells, thereby preventing their apoptosis (Hatzold and Conradt, 2008; Wei et al., 2020) However, the mechanism underlying the asymmetrical activation of *egl-1* in other ACDs remains elusive.

In this study, we demonstrate the enrichment of the NuRD complex in cells that are predetermined to survive, and its role in suppressing the EGL-1-CED-9-CED-4-CED-3 apoptotic pathway through repression of the *egl-1* gene. Furthermore, we report the interaction between the NuRD complex and the V-ATPase complex, and reveal that during asymmetric cell division, the localization of the V-ATPase complex and cytoplasmic pH are asymmetrical, thereby contributing to the polarized segregation of the NuRD complex.

## Results

### NuRD asymmetric segregation during neuroblast ACDs

In order to gain insights into the molecular distinctions between apoptotic and surviving cells in *C. elegans* embryos, we employed the SPLiT single-cell RNA sequencing (scRNA-seq) platform to compare their transcriptomes (Figure S1A-S1C) (Rosenberg et al., 2018). The expression of the critical somatic apoptosis-inducing gene *egl-1* was used to distinguish the apoptotic and surviving cells (Conradt et al., 2016). Intriguingly, we identified transcripts encoding subunits of the nucleosome remodeling and deacetylase (NuRD; also known as Mi-2) complex in cells where *egl-1* expression was indiscernible (Figure S1D-S1F, Supplementary Table 1). The NuRD complex represents an evolutionarily conserved protein complex associated with chromatin that couples the activities of chromatin-remodeling ATPases with histone deacetylases (Figure 1B) (Bracken et al., 2019; Lai and Wade, 2011). NuRD-mediated alterations in chromatin structure are vital for appropriate transcriptional regulation during cell fate determination and lineage commitment (Bracken et al., 2019; Lai and Wade, 2011), suggesting that NuRD may serve as a candidate epigenetic factor to specify cell fate between survival and death.

To investigate whether NuRD components are distributed asymmetrically between apoptotic and surviving daughter cells, we overexpressed GFP-tagged NuRD subunits within the *C. elegans* Q neuroblast lineages. Q.a neuroblast undergoes ACD to generate a large surviving Q.ap cell and a small apoptotic Q.aa cell (Figure 1A). We found that GFP-tagged NuRD subunits, including the histone deacetylase HDA-1, the histone binding protein LIN-53, the nucleosome remodeling factor CHD-3, and the MEP-1 protein were enriched asymmetrically in the surviving QR.ap cell during QR.a division (Figure S2, Supplementary Movie S1 and S2). Notably, NuRD asymmetric distribution occurred within several minutes after metaphase, while GFP protein translation and chromophore maturation require approximately 30 min in Q neuroblast (Ou and Vale, 2009), suggesting that the asymmetric enrichment of NuRD is likely the result of protein redistribution.

To investigate the dynamic distribution of endogenous NuRD during ACD, we generated a GFP knock-in (KI) strain for HDA-1 and an mNeonGreen (NG, green fluorescence) KI line for LIN-53 using CRISPR-Cas9 (Figure 1C). In QR.a cells, nuclear HDA-1 and LIN-53 were released into the cytoplasm and evenly distributed until metaphase (Figure 1D, Supplementary Movie S3 and S4). At anaphase, HDA-1 and LIN-53 became enriched in the posterior part of QR.a but became less detectable in the anterior part (Figure 1D, S3A-B, Supplementary Movie S3 and S4). We quantified the fluorescence intensity ratio between the posterior and anterior chromatids of QR.a, and between QR.ap and QR.aa nuclei, and found that nuclear HDA-1 or LIN-53 asymmetry gradually increased from 1.1-fold at anaphase onset to 1.5 or 1.8-fold upon completion of cytokinesis, respectively (Figure 1D-E). We also measured the ratios of fluorescence intensities between the posterior and anterior halves of QR.a, and between QR.ap and QR.aa (Figure S4A, see Materials and Methods). NuRD asymmetry became evident at ∼4 minutes and reached a plateau at ∼6 minutes after the anaphase onset (Figure 1D, 1F, S3A and S4B). QR.a spent ∼6 minutes from anaphase to the completion of cytokinesis (Chai et al., 2012; Ou et al., 2010), suggesting that QR.a cell establishes NuRD asymmetry during ACD.

Similar to QR.a, the dynamic and progressively developing NuRD asymmetry was also observed in the QL.a, QR.p and QL.p cell lineages, which generate apoptotic cells (Figure 1G, S3B and S4B). In contrast, an even distribution of HDA-1 and LIN-53 was observed in two surviving daughters of QR, QL, QL.pa and QR.pa cells, which generate two viable siblings (Figure 1G and S3B). To investigate whether other epigenetic factors are also asymmetrically segregated during Q cell ACDs, we tagged the MYST family histone acetyltransferase (MYS-1 in *C. elegans*) with GFP. We found that MYS-1::GFP was symmetrically segregated into the apoptotic and surviving daughter cells (Figure 1D-F, S3A and S4B, Supplementary Movie S5), indicating the specificity of the polarized NuRD partition. The total fluorescence of HDA-1, LIN-53, and MYS-1 remained constant during ACDs, suggesting that protein redistribution may establish NuRD asymmetry (Figure S4C). Inhibition of a PAR-1 family kinase PIG-1 led to symmetric Q cell divisions, producing extra neurons derived from some of their apoptotic daughters (Cordes et al., 2006). In *pig-1 (gm344)* mutants, HDA-1 and LIN-53 were evenly partitioned during ACDs (Figure S5, Supplementary Movie S6 and S7), indicating that asymmetric NuRD segregation depends on the PIG-1 kinase.

### NuRD asymmetric segregation in embryonic cell lineages

Given that a significant portion of somatic apoptotic events in *C. elegans*-113 out of 131-occur during embryonic development (Sulston et al., 1983a), we employed live imaging and automated cell lineage tracing algorithms (Du et al., 2014) to monitor the asymmetry of NuRD distribution in embryos from the early two- or four-cell stage up to the 350-cell stage. Our analysis indicated that in 15 out of 17 embryonic ACDs that produce apoptotic daughter cells, the surviving daughter cells showed enrichment of HDA-1 and LIN-53, with an average enrichment ratio exceeding 1.5-fold (Figure 2A and Supplementary Table 2). Notably, this enrichment reached statistical significance in 6 out of 17 embryonic ACDs (Figure 2B). Intriguingly, the MSpppaa cell lineage’s two daughter cells received comparable levels of NuRD, yet its apoptotic daughter cell completes apoptosis approximately 400 minutes post-division (Hsieh et al., 2012; Sulston et al., 1983b), supporting the hypothesis that lower NuRD concentrations might promote apoptosis. Conversely, the apoptotic daughter cell of ABaraaaap completes apoptosis within 35 minutes (Sulston et al., 1983a), despite receiving a similar amount of NuRD as its non-apoptotic sibling. Therefore, while the regulation of apoptosis in certain embryonic cells appears to be governed by NuRD-independent mechanisms, NuRD asymmetric segregation is evident in several embryonic cell lineages.

**Figure 2:**
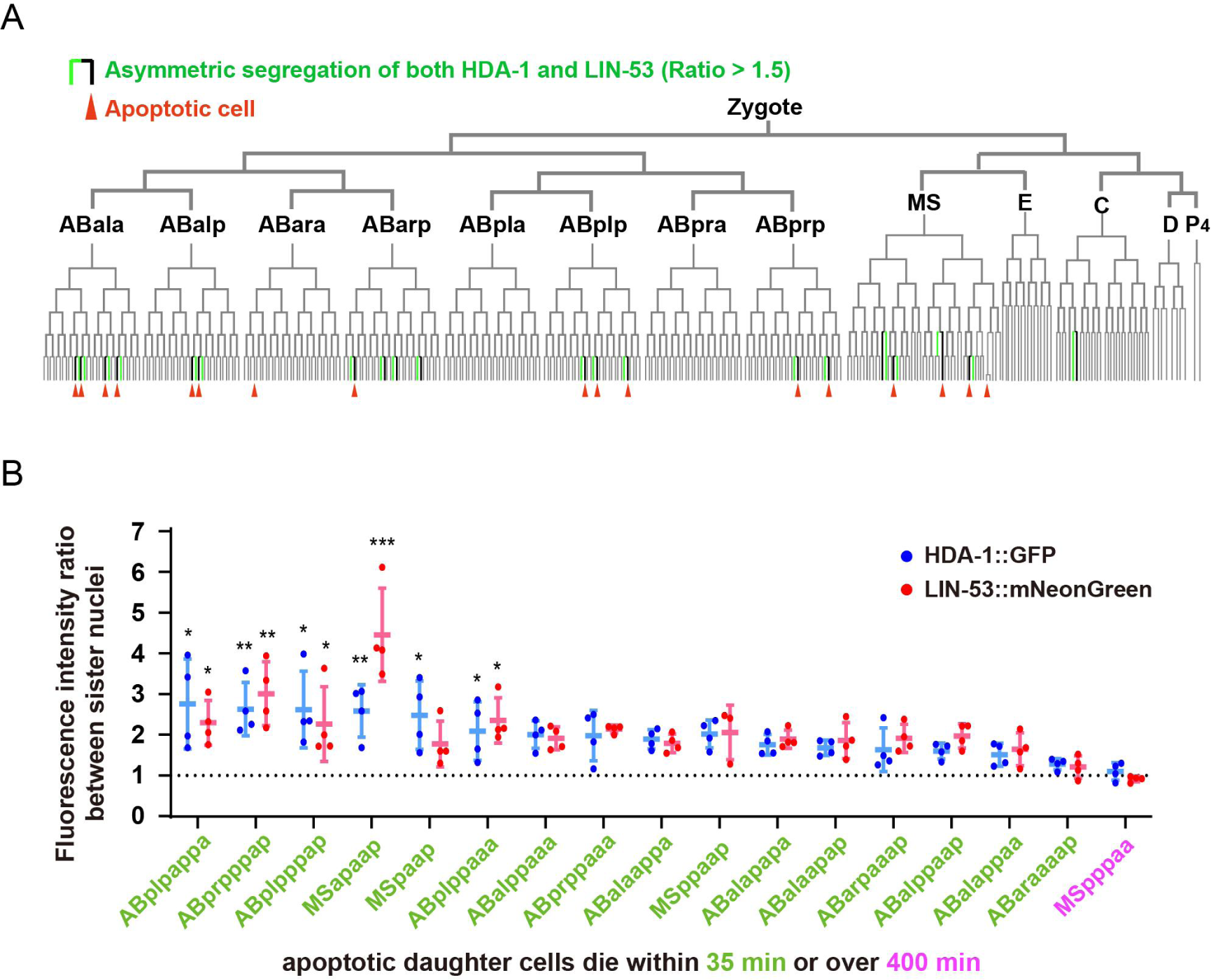
NuRD Asymmetry in *C. elegans* embryonic cell lineages. (A) The tree visualization depicts the segregation of HDA-1::GFP and LIN-53::mNeonGreen between sister cells during embryonic development. In this tree structure, vertical lines represent cells and horizontal lines denote cell divisions. Green vertical lines highlight cells with higher nuclear HDA-1::GFP and LIN-53::mNeonGreen fluorescence intensity than their apoptotic sister cells (average fluorescence intensity ratio between sister cell nuclei >1.5). Red arrowheads point to apoptotic cells. The placement of cells within the tree follows the Sulston nomenclature. See also Supplementary Table 2. (B) Quantifications of HDA-1::GFP and LIN-53::mNeonGreen fluorescence intensity ratios between nuclei of live daughter cells and their apoptotic sister cells. The lineage names of 17 cells that divide to produce apoptotic daughter cells are shown below the X-axis. Data are shown as mean ± SD. N = 3–4. The Dunn’s multiple comparisons test was used to assess statistical significance, with MSpppaa (magenta), whose apoptotic daughter cell completes apoptosis over 400 min after birth, as a control. *p < 0.05, **p < 0.01, ***p < 0.001.

### Loss of the deacetylation activity of NuRD causes ectopic apoptosis

To investigate the role of NuRD in determining cell fate, we reduced the deacetylation activity of NuRD using RNA-mediated interference (RNAi). The *hda-1* and *lin-53* RNAi efficacy was confirmed by the reduced green fluorescence in the germlines of HDA-1::GFP and LIN-53::mNeonGreen KI animals, as well as the increased acetylation level of histone H3K27 (Figure S6). As reported before, RNAi of either *hda-1* or *lin-53* caused embryonic lethality (Shi and Mello, 1998). RNAi of *lin-53* led to embryos arrested at gastrulation stage before the generation of apoptotic cells, preventing us from analyzing its roles in somatic apoptosis. In contrast, *hda-1* RNAi embryos arrested between the late gastrulation stage and bean stage, allowing us to tracked some of apoptotic events. To quantify apoptotic events, we used a secreted Annexin V (sAnxV::GFP) sensor to label apoptotic cells with externalized phosphatidylserine (exPS) on the surface of the plasma membrane in *ced-1(e1735)* mutant embryos which inhibits engulfment of apoptotic cells (Conradt et al., 2016; Mapes et al., 2012; Zhou et al., 2001). The sAnxV::GFP labeled an average of 10 cells in *ced-1(e1735)* mutant embryos treated with control RNAi and an average of 14 cells in *ced-1(e1735)*; *hda-1* RNAi embryos (Figure 3A-B). To test whether the upregulated apoptosis in *hda-1* RNAi embryos depends on the canonic apoptosis pathway, we introduced cell death-deficient mutations *ced-3/* Caspase *(n2433), ced-4/* APAF-1-like *(n1162), ced-9/* BCL-2-like *(n1950),* and *egl-1/* BH-3 only protein *(n1084n3082)* into *ced-1* mutant, respectively. RNAi of *hda-1* in these embryos only generated 0-2 exPS-positive cells (Figure 3A-B and S7A), indicating that the ectopic increase of apoptosis by HDA-1 inhibition requires the EGL-1-CED-9-CED-4-CED-3 pathway. Considering the pleiotropic phenotypes caused by loss of HDA-1, we cannot exclude the possibility that ectopic cell death might result from global changes in development, even though HDA-1 may directly contribute to the life-versus-death fate determination.

**Figure 3:**
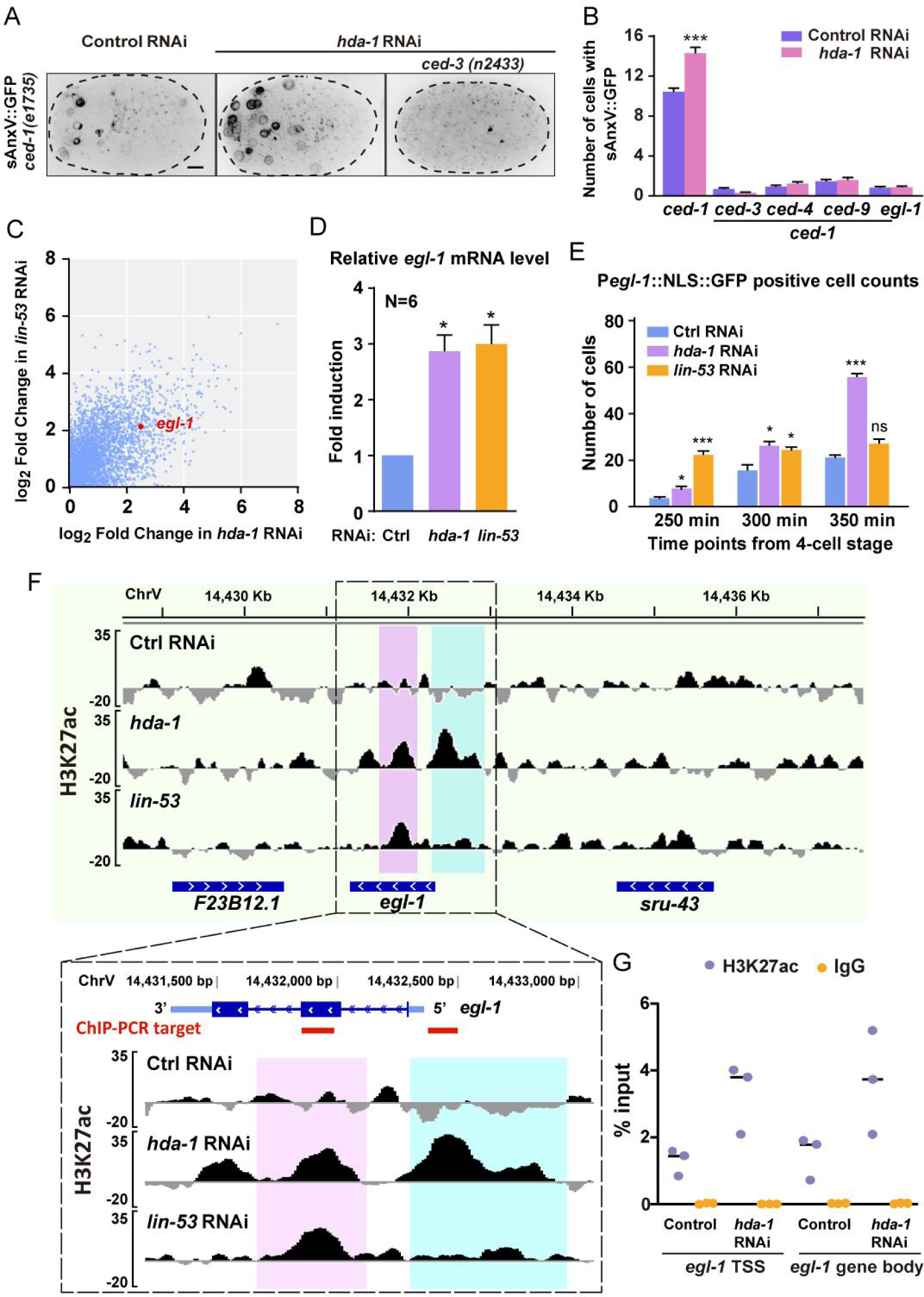
RNAi of *hda-1* induces ectopic apoptosis and increases H3K27 acetylation of the *egl-1* gene. (A) Representative inverted fluorescence images show P*hsp*::sAnxV::GFP from *ced-1(e1735)* and *ced-1(e1735); ced-3(n2433)* embryos between late gastrulation and bean stage, treated with control RNAi or *hda-1* RNAi. Scale bars, 5 μm. (B) Quantifications of cell corpse number in the embryos of indicated genotypes. Data are presented as mean ± SEM. N = 39–60. Statistical significance is determined by Student’s *t* test. ***p < 0.001. (C) Scatter plot of the increased gene expression in both *hda-1* RNAi and *lin-53* RNAi animals. The *egl-1* gene is marked in red. See also Supplementary Table 3. (D) Quantitative real-time PCR (RT-PCR) measurement of *egl-1* mRNA levels in the control, *hda-1* or *lin-53* RNAi embryos. Fold induction was calculated relative to levels in control RNAi embryos. Data of six biological replicates are presented as mean ± SEM. Statistical significance is determined by the Wilcoxon test as 1 as the theoretical median. *p < 0.05. (E) Quantification of the number of cells expressing the P*egl-1*::NLS::GFP reporter in embryos treated with control, *hda-1* or *lin-53* RNAi. Data are presented as mean ± SEM. N = 8–9. Statistical significance is determined by the Dunn’s multiple comparisons test. *p < 0.05, ***p < 0.001, ns: not significant. (F) Normalized ChIP-Seq signal profiles of the H3K27 acetylation level in the control, *hda-1* or *lin-53* RNAi embryos at the *egl-1* locus. The Y-axis shows the average sequencing coverage of bins per million reads of three biological replicates normalized to the input control sample. Ectopic H3K27ac enrichments at the *egl-1* locus of *hda-1* RNAi and *lin-53* RNAi embryos are highlighted in magenta and cyan. (G) ChIP-qPCR analyses using H3K27ac antibody or IgG (the negative control) at selected elements of *egl-1* indicated in (F) by red lines. Results are shown as the percentage of input DNA. Data of three biological replicates are presented.

### NuRD RNAi upregulates the *egl-1* expression by increasing its H3K27 acetylation

To understand how NuRD regulates apoptotic cell fates, we performed RNA-Seq analyses on WT, *hda-1* RNAi and *lin-53* RNAi embryos. In both RNAi conditions, we observed 287 or 575 genes that were upregulated or downregulated (|log_2_FD| > 2 and FDR < 0.05) (Supplementary Table 3). Notably, we found that the apoptosis-inducing gene *egl-1* was among the upregulated genes (Figure 3C). Our quantitative RT-PCR results further confirmed that *egl-1* transcripts were more abundant in *hda-1* RNAi and *lin-53* RNAi embryos than in control RNAi embryos (Figure 3D). We also used a P*egl-1*::NLS::GFP transcriptional reporter to examine the *egl-1* transcription activity during embryonic development, which allowed us to detect *egl-1* expression while avoiding the adverse effects of *egl-1* overexpression on apoptosis. We found that RNAi of *hda-1* or *lin-53* caused an increased number of P*egl-1*::NLS::GFP positive cells (Figure 3E and S7B). These results indicate that the loss of NuRD abnormally activates the *egl-1* gene expression.

Next, we investigated whether the loss of histone deacetylase activity in NuRD increased the H3K27ac levels at the *egl-1* locus, as the histone H3K27ac levels are associated with transcription activation (Heinz et al., 2015). To test this, we performed ChIP-Seq experiments with the anti-H3K27ac antibody and found that the transcription start site (TSS) and the gene body region of *egl-1* had higher levels of H3K27ac in *hda-1* or *lin-53* RNAi embryos compared to WT, while the H3K27ac levels of genes adjacent to *egl-1* showed no significant changes (Figure 3F). This was supported by ChIP-qPCR in *hda-1* RNAi embryos (Figure 3G). Therefore, inhibiting NuRD increased the H3K27ac levels at the *egl-1* locus, leading to its upregulation and subsequent activation of apoptosis.

### V-ATPase regulates asymmetric segregation of NuRD during somatic ACDs

To investigate how NuRD is asymmetrically segregated during ACD, we used affinity purification with an anti-GFP antibody and mass spectrometry (MS) to isolate NuRD binding partners from the lysate of HDA-1::GFP KI animals. Our analysis revealed all the previously known NuRD subunits from HDA-1::GFP KI animals (Figure S8, Supplementary Table 4) (Bracken et al., 2019; Lai and Wade, 2011), validating our experimental method. Interestingly, our co-immunoprecipitation and MS with HDA-1::GFP identified 12 subunits of the vacuolar-type H^+^-ATPase (V-ATPase) (Figure 4A-B), consistent with a previous report that used FLAG-tagged LIN-53 to identify 4 V-ATPase subunits (Muthel et al., 2019). We further confirmed the binding of HDA-1 to V-ATPase subunits in *C. elegans* by co-immunoprecipitation and western blot analysis with anti-V1A/VHA-13 antibodies (Figure 4C). V-ATPase is a proton pump that comprises a transmembrane domain (V0) responsible for proton transport across membranes and a peripheral ATP-hydrolytic domain (V1) that catalyzes ATP hydrolysis (Figure 4B). V-ATPases play a critical role in intracellular pH homeostasis and regulate numerous physiological processes, including apoptosis (Abbas et al., 2020; Chen et al., 2022; Ernstrom et al., 2012; Vasanthakumar and Rubinstein, 2020). However, the role of V-ATPase in the asymmetric segregation of epigenetic factors during ACD remains unknown.

**Figure 4:**
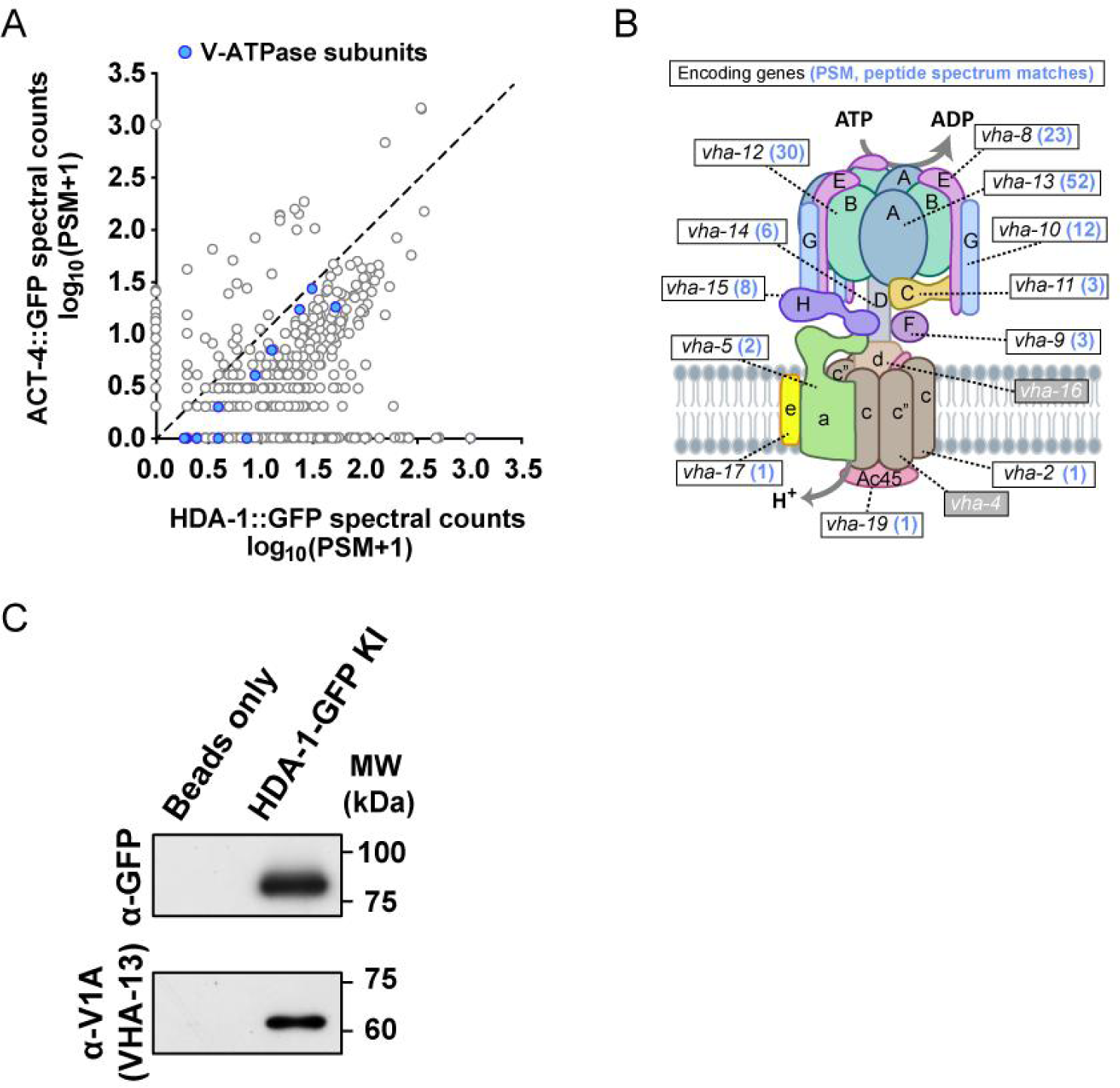
HAD-1 interacts with subunits of V-ATPase. (A) The plot compares counts of proteins co-precipitated with HDA-1::GFP with those with the control ACT-4 (actin)::GFP. The PSM (Peptide-Spectrum Match) is the number of identified peptide spectra matched for the protein. Blue dots represent the subunits of V-ATPase. See also Supplementary Table 4. (B) Schematic model of the V-ATPase complex. The known worm subunits are indicated. The mean PSM of the encoded protein from two co-IP and MS repeats is shown in blue after the gene name. (C) Western blot (WB) showing co-immunoprecipitation (co-IP) of V-ATPase V1 domain A subunit (V1A) with the HDA-1 from worm lysates. Assay was performed using three biological replicates. Three independent biological replicates of the experiment were conducted with similar results

To investigate the role of V-ATPase in NuRD asymmetric segregation in Q neuroblast, we used a pharmacologic inhibitor, bafilomycin A1 (BafA1), to inhibit V-ATPase proton pumping activity (Furuchi et al., 1993; Wang et al., 2021; Yoshimori et al., 1991). To assess the inhibitory effect of BafA1 on the proton translocation activity of V-ATPase in Q neuroblast, we monitored the cytosolic pH dynamics in dividing QR.a using the super-ecliptic pHluorin. The pHluorin is a pH-sensitive GFP reporter whose fluorescence can be quenched by the acidic pH (Miesenbock et al., 1998; Sankaranarayanan et al., 2000). Although BafA1-mediated disruption of lysosomal pH homeostasis is recognized to elicit a wide array of intracellular abnormalities, we found no evidence of such pleiotropic effects at the organismal level with the dosage and duration of treatment employed in this study. In DMSO-treated animals, the pHlourin fluorescence intensity remained constant in the posterior portion that forms QR.ap but was significantly reduced in the anterior portion that forms QR.aa, indicating that the cytoplasm of the future apoptotic cell became more acidic (Figure 5A-B and Supplementary Movie S8). This observation is consistent with the recognition that cytosolic acidification is a common feature of both death receptor-mediated and mitochondria-dependent apoptosis (Lagadic-Gossmann et al., 2004; Matsuyama et al., 2000; Matsuyama and Reed, 2000). Notably, BafA1 treatment reduced the pHluorin fluorescence intensity ratio between QR.ap and QR.aa from 1.6-fold to 1.3-fold (Figure 5A-B and Supplementary Movie S8), suggesting that BafA1 may disrupt the cytosolic pH asymmetry in dividing QR.a cells by inhibiting V-ATPase activity, although we cannot exclude the possibility that the changes in fluorescence could be due to changes in the amount of pHluorin protein.

**Figure 5:**
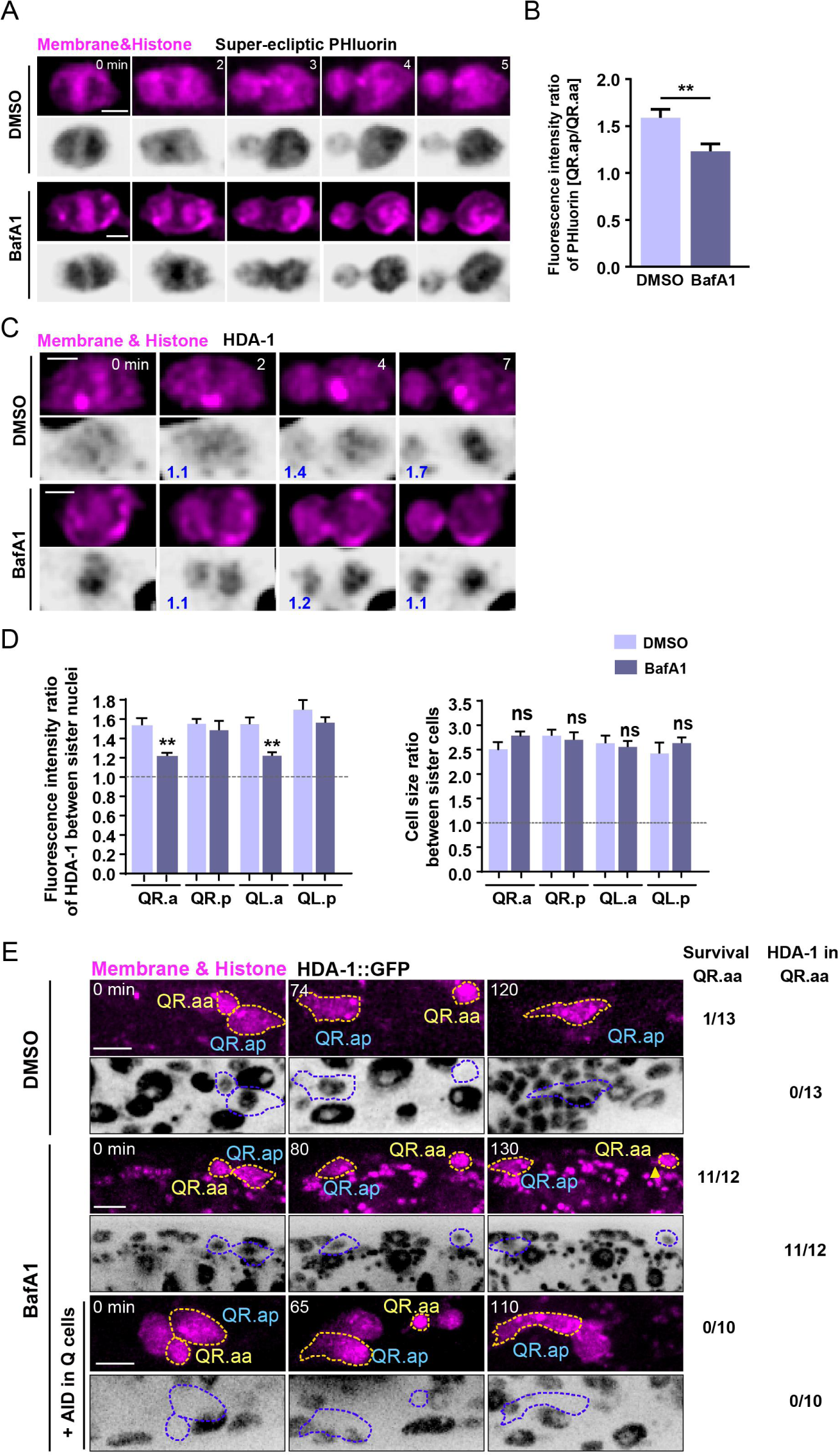
V-ATPase regulates NuRD asymmetric segregation and cell fates. (A) Dynamics of the cytosolic pH indicated by super-ecliptic pHluorin during QR.a division in DMSO- or BafA1-treated animals. In each panel, the top row shows mCherry-tagged plasma membrane and histone, and the bottom row shows inverted fluorescence images of super-ecliptic pHluorin. The anterior of the cell is on the left. Time 0 min is the onset of anaphase. Scale bar, 2 µm. See also Supplementary Movie S8. (B) The super-ecliptic pHluorin fluorescence intensity ratio between QR.ap and QR.aa in DMSO control or BafA1-treated animals. Data are presented as mean ± SEM. N = 11–15. Statistical significance is determined by Student’s *t* test. **p < 0.01. (C) Images of HDA-1::GFP distribution during QR.a cell division in DMSO- or BafA1-treated animals. In each panel, the top row shows mCherry-tagged plasma membrane and histone, and the bottom row shows inverted fluorescence images of HDA-1::GFP. The GFP fluorescence intensity ratios between the posterior and anterior chromatids, and between QR.ap and QR.aa nuclei are shown in blue at the lower left corner of inverted fluorescence images. Anterior of the cell is left. Scale bar: 2 µm. See also Supplementary Movie S9. (D) Quantification of HDA-1::GFP fluorescence intensity ratios between the large and small daughter cell nuclei and the cell size ratios between the large and small daughters. The names of mother cells are shown on the X-axis. Data are presented as mean ± SEM. N = 9–12. Statistical significance is determined by Student’s *t* test. **p < 0.01, ns: not significant. (E) Representative images showing fates of QR.aa after DMSO (top), BafA1 (middle) and BafA1 plus AID treatment (bottom). The Q cell plasma membrane and chromosome are labeled by mCherry. HDA-1::GFP is shown as inverted fluorescence images. Yellow arrow head shows a short neurite-like outgrowth of QR.aa in BafA1 treated larvae. Frequencies of QR.aa survival and HDA-1 maintenance are showed on the right. Time 0 min is the birth of QR.aa. Scale bar: 5 µm.

We observed that BafA1 treatment resulted in the symmetric segregation of HDA-1 during QR.a division (Figure 5C-D and Supplementary Movie S9). Notably, neither DMSO nor BafA1 treatment affected the asymmetry in daughter cell size (Figure 5C-D and Supplementary Movie S9), suggesting that intracellular acidification, regulated by V-ATPase activity, specifically affects the NuRD asymmetric segregation without affecting the asymmetry in daughter cell size. We also investigated whether the small QR.aa cell carrying ectopic NuRD could escape apoptosis. In DMSO-treated animals, QR.aa cells underwent apoptosis, and 12 out of 13 QR.aa cells were engulfed and degraded by a neighboring epithelial cell within 120 min after birth. However, in BafA1-treated animals, QR.aa inherited similar levels of HDA-1::GFP as its sister cell, and 11 out of 12 QR.aa cells carrying ectopic NuRD survived for over 120 min and formed a short neurite-like outgrowth (Figure 5E). To confirm whether the long-lived QR.aa was a consequence of the ectopic gain of NuRD, we depleted HDA-1 within Q cell lineages using the auxin-inducible protein degradation (AID) system (Zhang et al., 2015). By introducing the degron sequence into the *hda-1* locus and expressing the TIR1 F-box protein under the Q cell-specific *egl-17* promoter, we showed that HDA-1::GFP fluorescence in Q cells was substantially decreased under auxin treatment (Figure 5E). Administration of auxin restored QR.aa apoptosis in BafA1-treated animals (Figure 5E). These results suggest that V-ATPase activity-dependent NuRD asymmetric segregation contributes to the specification of the live-death fate.

To understand how V-ATPase regulates NuRD asymmetric segregation, we generated a transgenic strain expressing wrmScarlet-tagged VHA-17, which is the e subunit of the V0 domain. Using this strain, we were able to examine the dynamic distribution of V-ATPase during QR.a cell division. Like HDA-1, V-ATPase was also asymmetrically enriched in the surviving QR.ap portion (Figure 6A-B and Supplementary Movie S10). This observation suggests that V-ATPase and NuRD may co-segregate asymmetrically during ACD. We also found that BafA1 treatment disrupted V-ATPase asymmetric distribution (Figure 6A-B and Supplementary Movie S10), indicating the importance of the proton-pumping activity of V-ATPase in its asymmetric segregation. Therefore, our results suggest that V-ATPase may facilitate the asymmetric distribution of NuRD through its proton pumping activity.

**Figure 6:**
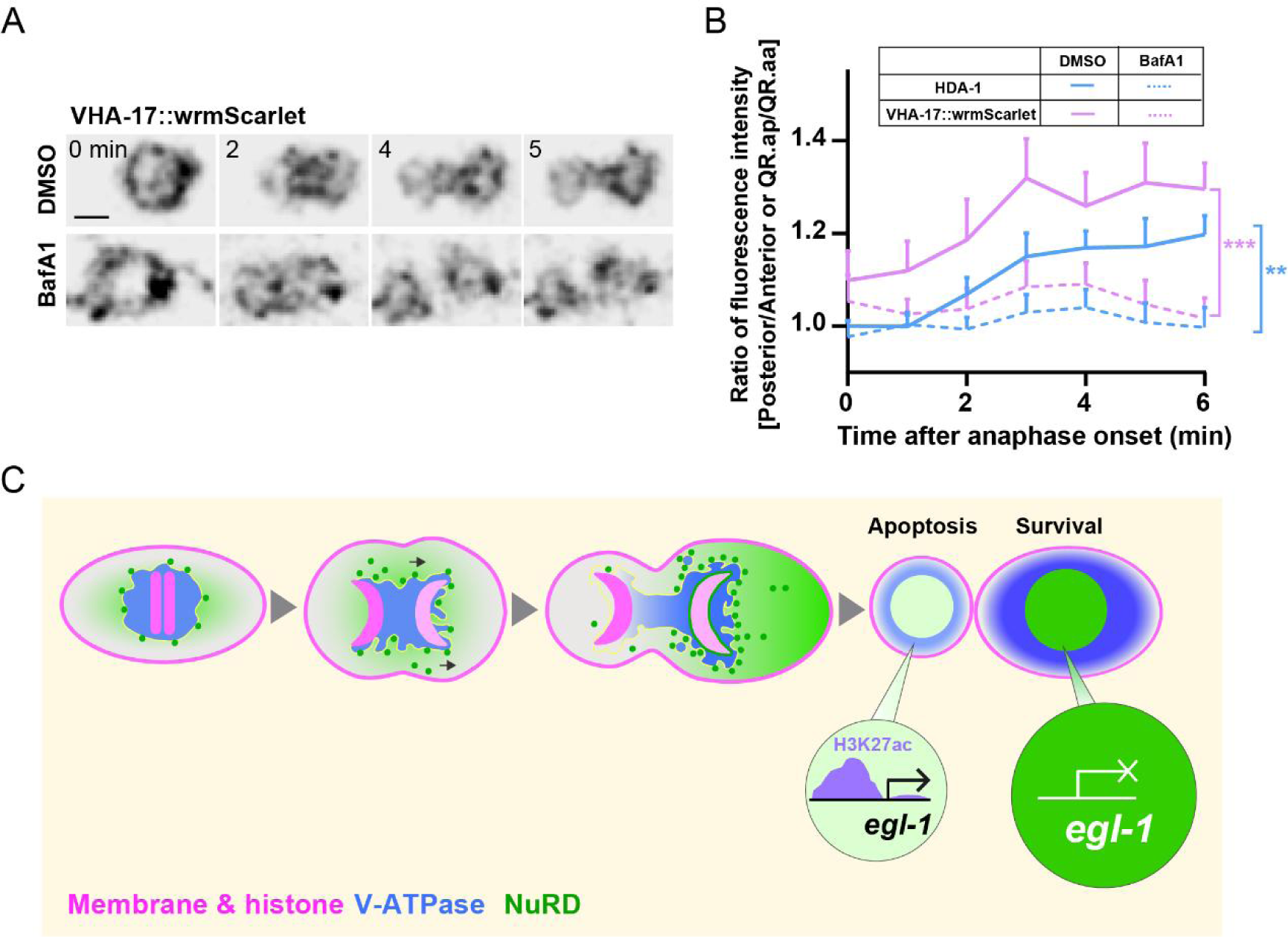
V-ATPase distribution during ACDs and a model. (A) Dynamics of VHA-17::wrmScarlet during QR.a cell division in DMSO- or BafA1-treated animals. VHA-17::wrmScarlet fluorescence is shown as inverted fluorescence images. Time 0 min is the onset of anaphase. Scale bar: 2 µm. See also Supplementary Movie S10. (B) Quantification of VHA-17::wrmScarlet (magenta) and HDA-1::GFP (blue) fluorescence intensity ratios between the posterior and anterior half of QR.a or between QR.ap and QR.aa. Data are presented as mean ± SEM. N = 10–14. Student’s *t* test was used to compare the intensity ratio difference of VHA-17 and HDA-1 between DMSO- and BafA1-treated cells at 6 min after anaphase. **p < 0.01, ***p < 0.001. (C) A proposed model. Asymmetric segregation of V-ATPase mediated the enrichment of its associated NuRD in the large daughter cell, where high level of NuRD deacetylates *egl-1* and suppressed its expression.

## Discussion

Our findings indicate that the asymmetric segregation of the NuRD complex during ACD is regulated in a V-ATPase-dependent manner, and is critical for the differential expression of apoptosis activator *egl-1* and the life-versus-death fate decision (Figure 6C). Specifically, we have provided evidence that the reduced level of NuRD in the apoptotic daughter cell leads to an increased level of H3K27ac, an epigenetic modification known to be associated with active gene expression, at the *egl-1* locus, resulting in the upregulation of *egl-1* expression and subsequent induction of apoptosis. Based on our results, we propose a dual-function model of V-ATPase in NuRD asymmetric segregation: (i) V-ATPase interacts with NuRD, both of which are asymmetrically segregated into the surviving cell; (ii) the asymmetrical activity of V-ATPase generates a more acidic cytoplasmic environment in the future apoptotic cell portion compared to the surviving portion, which might contribute to the asymmetric segregation of V-ATPase and NuRD. This model sheds light on the mechanisms underlying the regulation of apoptosis during ACD, and opens up new avenues for further investigation into the intricate interplay between V-ATPase, NuRD, and epigenetic modifications in the context of cell fate decisions.

The transcript asymmetry detected by scRNA-seq may not correspond to the protein asymmetry detected by microscopic imaging. Our scRNA-seq data shows that 6487 out of 8624 genes were not detected in *egl-1*-positive cells, the putative apoptotic cells. Cells that are *egl-1* positive may be undergoing apoptosis, rendering the asymmetry of NuRD complex transcripts insignificant in inferring protein asymmetry. Thus, the observed transcript asymmetry of the NuRD subunits between live and dead cells may be coincidental with NuRD protein asymmetry during asymmetric neuroblast division, rather than serving as a regulatory mechanism.

The intrinsic mechanisms governing binary cell fate decisions involve asymmetric cortical localization of cell fate determinants, polarized partitioning of RNA species, unequal segregation of organelles, and biased distribution of damaged proteins or protein aggregates (Sunchu and Cabernard, 2020; Venkei and Yamashita, 2018). Despite this, little is known about the asymmetric segregation of epigenetic modification enzymes during ACDs, nor the functions of polarized V-ATPase distribution and cytoplasmic proton asymmetry during this process. Our findings provide new insights into the asymmetric segregation of cell-intrinsic factors and the previously unrecognized roles of V-ATPase and cytosolic acidification in this process.

Two mechanisms have been proposed to explain the V-ATPase-dependent asymmetric segregation of NuRD. Firstly, a polarized intracellular transportation system could selectively deliver NuRD-V-ATPase-containing organelles to the surviving daughter cell. Although V-ATPase is primarily known for its localization and function in the late endosome and lysosome, recent evidence suggests that V-ATPase subunits are synthesized and assembled on the endoplasmic reticulum (ER) (Abbas et al., 2020; Graham et al., 2003; Wang et al., 2020). Our findings demonstrate that V-ATPase colocalizes with the ER marker during ACDs, suggesting that NuRD-V-ATPase may be delivered as cargo on the ER or ER-derived vesicles (Figure S9). As reported in previous studies, a polarized microtubule transportation system, governed by specific motor proteins and the microtubule tracks, is responsible for the asymmetric segregation of signaling endosomes during ACDs of *Drosophila* sensory organ precursors (Derivery et al., 2015). It is possible that a similar biased transportation system plays a role in the asymmetric segregation of NuRD-V-ATPase-containing organelles.

Secondly, the surviving daughter cell chromatids may have a greater ability to recruit NuRD than their apoptotic sister. This could be due to a higher number of yet unknown NuRD recruiting factors on chromatids or specific posttranslational modifications of chromosomal proteins that enhance the chromosomal recruitment of NuRD. Despite having an identical DNA sequence, sister chromatids can be distinguished by chromosomal proteins or modifications (Ranjan et al., 2019; Tran et al., 2012), which likely provides the specific molecular cues for asymmetric NuRD recruitment. The NuRD asymmetry on chromosomes (1.5∼1.8-fold between QR.ap and QR.aa nuclei) is greater than that between the two daughter cells (approximately 1.3-fold) (Figure 1D-1F), suggesting that polarized NuRD-V-ATPase transportation and distinct NuRD affinity from sister chromatids may act in concert to establish NuRD asymmetry.

The observation of asymmetric segregation of the NuRD complex during cell divisions suggests that the mother cell may play a role in initiating the fate specification of its daughter cells. This notion is supported by studies conducted on *Drosophila* male germline stem cells (GSCs) which demonstrated that the old and new histone H3-H4 or unequal amounts of histone H3 variant CENP-A were incorporated into sister chromatids prior to cell division (Ranjan et al., 2019; Tran et al., 2012). Although these patterns were not observed in cell types other than GSCs, asymmetric NuRD segregation was found to be common in *C. elegans* (Figure 1D-G, Figure 2A, and Figure S3). In 395 tracked embryonic cell divisions that generated two surviving daughter cells, NuRD was equally distributed in 390 cases, but in five cases, significant differences in HDA-1 and LIN-53 levels were observed between siblings (Figure 2A, Supplementary Table 2). This observation suggests that biased NuRD segregation may contribute to binary cell fate decisions beyond apoptosis. In the embryonic cerebral cortex of mammals, approximately 30% of newborn cells undergo programmed cell death (Voiculescu et al., 2000), which is a significantly higher rate than observed in *C*. *elegans* where only 12% of somatically born cells undergo apoptosis (131 apoptotic cells among 1090 somatically born cells). Thus, it is worth exploring whether brain stem cell divisions employ asymmetric NuRD segregation to specify daughter cell fates. Further studies on polarity establishment hold promise in elucidating the mechanisms underlying the asymmetric inheritance of epigenetic information.

## Materials and Methods

### Worm strains and culture

*C. elegans* were maintained as described in the standard protocol (Brenner, 1974). All strains were cultivated at 20 °C on nematode growth medium (NGM) agar plates seeded with *Escherichia coli* OP50 or HT115 (feeding RNAi assay). The wild-type strain was Bristol N2. Some strains were provided by the *Caenorhabditis* Genetics Center (CGC), funded by the NIH Office of Research Infrastructure Programs (P40 OD010440). The strains used in this study were listed in Extended Data Table 1.

### Molecular biology

We performed genome editing experiments in *C. elegans* following the established methods (Dickinson et al., 2013; Friedland et al., 2013). We used the CRISPR design tool (https://zlab.bio/guide-design-resources) to select the target sites. The sgRNA sequences (Extended Data Table 2) were inserted into the pDD162 vector (Addgene #47549) by linearizing the vector with 15 base pairs (bp) overlapped primers using Phusion high-fidelity DNA polymerase (New England Biolabs, cat. no. MO531L). PCR products were digested using Dpn I (Takara, cat. no. 1235A) for 2 hours at 37 °C and transformed into Trans5 α bacterial chemically competent cells (TransGen Biotech, Cat no. CD201-01). The linearized PCR products with 15 bp overlapping ends were cyclized to generate plasmids by spontaneous recombination in bacteria. Homology recombination (HR) templates were constructed by cloning the 1.5 kb upstream and downstream homology arms into the pPD95.77 plasmid using the In-Fusion Advantage PCR cloning kit (Clontech, Cat. #639621). Fluorescence tags were inserted into the constructs with a flexible linker before the stop codons. Synonymous mutations were introduced to Cas9 target sites to avoid the cleavage of the homologous repair template by Cas9. The plasmids were listed in Extended Data Table 3.

Constructs that express GFP-tagged NuRD components were generated using the PCR SOEing method(Hobert, 2002). N2 genomic sequences (1.5-2 kb promoter plus coding region) were linked with *gfp*::*unc-54* 3’ UTR DNA fragments. Constructs that express wrmScarlet-tagged V-ATPase components with *gfp*::*unc-54* 3’ UTR were generated using the In-Fusion Advantage PCR cloning kit (Clontech, Cat. #639621). The primers were listed in Extended Data Table 3.

### Genome editing and transgenesis

To generate a knock-in strain, we purified the sgRNA construct and the repair template plasmids with the PureLink Quick PCR purification Kit (Invitrogen, #K310001) and co-injected them into N2 animals with the pRF4 [*rol-6 (su1006)*] and P*odr-1::dsRed* selection markers. The F1 transgenic progenies were singled and screened by PCR and Sanger sequencing. Transgenic *C. elegans* was generated by the germline microinjection of DNA plasmids or PCR products into N2 with P*odr-1::gfp* or P*egl-17*::*myri*-*mCherry* and P*egl-17::mCherry::his-24* plasmids. We maintained at least two independent transgenic lines with a constant transmission rate (>50%). Concentrations of DNA constructs used for generating knock-in were 50 ng/µl, and for transgenesis were 20 ng/µl. All the strains, primers, plasmids, and PCR products were listed in Extended Data Table 1-3, respectively.

### Live-Cell Imaging and quantification

Live imaging of *C. elegans* embryos, larvae, or young adults followed our established protocols (Chai et al., 2012; Zhang et al., 2017). L1 larvae or young-adult worms were anesthetized with 0.1 mmol/L levamisole in M9 buffer, mounted on 3% (wt/vol) agarose pads maintained at 20°C. Our imaging system includes an Axio Observer Z1 microscope (Carl Zeiss MicroImaging) equipped with a Zeiss 100×/1.46 numerical aperture (NA) objective, an Andor iXon+ EM-CCD camera, and 488 and 561-nm lines of a Sapphire CW CDRH USB Laser System attached to a spinning disk confocal scan head (Yokogawa CSU-X1 Spinning Disk Unit). Images were acquired by μManager (https://www.micro-manager.org). Time-lapse images of Q cell divisions were acquired with an exposure time of 200 msec every 1 min. Recent advancements in optical and camera technologies permit the acquisition of Z-stacks without perturbing Q cell division or overall animal development. Z-stack images were acquired over a range of −1.6 to +1.6 μm from the focal plane, at intervals of 0.8 μm. The field-of-view spaned 160 μm × 160 μm, and the laser power, as measured at the optical fiber, was approximately 1 mW. ImageJ software (http://rsbweb.nih.gov/ij/) was used to perform image analysis and measurement. Image stacks were z-projected using the average projection for quantification and using the maximum projection for visual display. To follow the cell fate of QR.aa in WT, BafA1- or BafA1 and IAA-treated worms, time-lapse images were acquired with 300 msec for 561 nm and 100 msec for 488 nm exposure time at 5-minute intervals. To image P*hsp-16.41*::sAnxV::GFP positive cells in WT, mutant, or RNAi animals, we incubated the eggs at 33 °C for 30 min to induce the reporter expression 3 hours post-egg-laying. After we kept the eggs at 20 °C for 2 hours, we imaged them with an exposure time of 200 msec. To acquire time-lapse images of P*egl-1*::NLS::GFP positive cells during embryonic development, we dissected 2-4 cell-stage embryos from gravid adults and imaged them with an exposure time of 300 msec every 5 min.

The quantifications of cellular fluorescence intensity ratios in Q cell lineages were described in Figure S4A. We used the mCherry-labeled plasma membrane to circumscribe Q cells (Region of Interest). To determine the ratios of fluorescence intensities in the posterior to anterior half (P/A) of Q.a lineages or A/P of Q.p lineages, the cell in the mean intensity projection was divided into posterior and anterior halves. ImageJ software was used to measure the mean fluorescence intensities of two halves with background subtraction. The slide background’s mean fluorescence intensity was measured in a region devoid of worm bodies. The background-subtracted mean fluorescence intensities of the two halves were divided to calculate the ratio. The same procedure was used to determine the fluorescence intensity ratios between two daughter cells. Total fluorescence intensity was the sum of the posterior and anterior fluorescence intensities or the sum of fluorescence intensities from two daughter cells (Figure S4A).

The ROIs for measuring the mean fluorescence intensities of nuclei were delineated as areas containing mCherry-tagged histone within each cell. The background-subtracted mean fluorescence intensities of the two nuclei were divided to calculate the nuclear intensity ratios.

### Feeding RNAi

RNAi bacteria were grown in LB media containing carbenicillin (50 µg/mL) and tetracycline hydrochloride (12.5 µg/mL) at 37 °C overnight. RNAi bacteria were then seeded on NGM plates supplemented with 50 µg/mL carbenicillin and 1 mM Isopropyl β-D-thiogalactopyranoside (IPTG). We grew the seeded RNAi plates for 2 days at room temperature, allowing double-stranded RNA expression. To score phenotypes during embryogenesis or meiosis, we transferred the synchronized young adult worms to the culture plate containing the induced *E. coli* HT115 strains expressing the control (luciferase) or specific gene-targeted dsRNAs. We incubated them for 16-24 hours at 20°C and then scored the phenotypes. As for RT-qPCR and RNA-Seq experiments, parental animals of RNAi were used for laying eggs, and the progeny was collected 5 hours post-egg-laying. In ChIP-Seq experiments, we used worms of mixed stages under RNAi treatment for 24-36 hours. RNAi bacterial strains were from the Vidal library.

### Immunoprecipitation and mass spectrometry

GFP transgenic or knock-in worms were raised on one hundred 90-mm NGM plates. Animals were collected and washed with M9 buffer three times. For each replicate, the lysate was made from 1∼2 ml packed worms with 3∼4 ml of 0.5-mm diameter glass beads using FastPrep-24 (MP Biomedicals) in lysis buffer [pH 7.4, 150 mM NaCl, 25 mM Tris-HCl, 10% glycerol, 1% NP-40, 1x cocktail of protease inhibitors from Roche (Complete, EDTA free), 40 mM NaF, 5 mM Na_3_VO_4_]. Worm lysates were then cleared by centrifugation at 14,000g for 30 min at 4 °C. For anti-GFP immunoprecipitation (IP), the supernatant was incubated with GFP-Trap A beads (Chromoteck, GTA20) for 1 hr. Beads were then washed three times with lysis buffer. To detect the interaction between the V-ATPase subunit and HDA-1, beads are boiled with 1X SDS-PAGE sample loading buffer for SDS–PAGE and western blot. Otherwise, proteins were eluted from beads with 300 μl 0.1 M glycine-HCl (pH 2.5) Add 15 μl 1.5 M Tris-HCl (pH 8.8) to the eluted protein, followed by precipitation using 100 μl trichloroacetic acid and re-dissolution in 60 μl 8 M Urea, 100 mM Tris-HCl (pH 8.5) for MS. MS Samples were then reduced in 5 mM TCEP, alkylated by 10 mM iodoacetamide, and diluted four-fold with 100 mM Tris-HCl (pH 8.5). The 0.2 μg trypsin was used to digest 10 μg protein in 1 mM CaCl_2_ and 20 mM methylamine overnight at 37 °C. The resultant peptides were desalted by ZipTip pipette tips (Merck Millipore). For liquid chromatography-tandem mass spectrometry analysis, peptides were separated by an EASY-nLCII integrated nano-HPLC system (Proxeon, Odense, Denmark) directly interfaced to a Thermo Scientific Q Exactive mass spectrometer (Thermo Fisher Scientific). Peptides were loaded on an analytical fused-silica capillary column (150 mm in length, 75 mm in internal diameter; Upchurch) packed with 5 mm, 300 Å C-18 resin (Varian). Mobile phase A consisted of 0.1% formic acid, while mobile phase B consisted of 0.1% formic acid and 100% acetonitrile. The Q Exactive mass spectrometer was operated through Xcalibur 2.1.2 software in data-dependent acquisition mode. A single full-scan mass spectrum was used in the orbitrap (400–1,800 m/z in mass range, 60,000 in resolution) followed by ten data-dependent tandem mass spectrometry scans at 27% normalized collision energy (HCD). The tandem mass spectrometry spectra results were searched against the *C. elegans* proteome database by the Proteome Discoverer (version PD1.4; Thermo Fisher Scientific).

### Generation of *C. elegans* embryonic cell suspensions for SPLiT-Seq

We prepared single cells from a *C. elegans* strain (*smIs89*) that expressed a P*egl-1*::NLS::GFP reporter using the established protocols with minor modifications(Bianchi and Driscoll, 2006; Zhang et al., 2011). Synchronized gravid adults were transferred to NGM plates to lay eggs for 1 hour, and about 50 μl eggs pellet were collected 3 hours post laid. We washed eggs three times with 1 ml sterile egg buffer [118 mM NaCl, 48 mM KCl, 3 mM CaCl_2_, 3 mM MgCl_2_, 5 mM HEPES (pH 7.2)]. We rinsed the pellet with 250 μl 1 U/ml Chitinase three times. Eggshells were digested with 250 μl of 1 U/ml chitinase for 45 min on a rotter. We stopped the reaction by adding 750 µl of L-15/FBS (L-15 medium containing 10% fetal bovine serum). Eggs were pelleted by centrifugation at 900 g for 3 min. The supernatant was displaced with 500 µl L-15/FBS. Embryos were dissociated by gently pipetting up and down 60 times using an insulin syringe (29 G needle). The dissociated cells were separated from undissociated cell clumps by filtering through a 5 µm mesh filter, and we washed the filter with 500 µl L-15/FBS. Cells were collected by centrifuging 1 ml filtered suspension at 3220 g for 3 min at 4 °C. We removed the supernatant and gently resuspended cells with 2 ml 1X PBS containing 5 μl SUPERaseIn RNase Inhibitor (Invitrogen) and 2.5 μl RiboLock RNase Inhibitor (Thermo Scientific).

### SPLiT-Seq library preparation and sequencing

We performed SPLiT-Seq as previously described (Rosenberg et al., 2018) with minor modifications. After cell fixation, centrifugations were performed at 3220 g for 3 min according to the *C. elegans* cell size. The RT primers were poly-dT and random hexamer mixture with UMIs (Unique Molecular Identifiers). All the barcodes used in this study were described in the published protocol(Rosenberg et al., 2018). The first 24 barcodes were used in the first round of barcoding, named “Round1_01” to “Round1_24”, and 48 barcodes were used in the third round of barcoding, called “Round3_49” to “Round3_96”. Each cell barcode came from three rounds of barcoding. TruePrep DNA Library Prep Kit V2 constructed the sequencing library for Illumina (Vazyme Biotech #TD502) according to the manufacturer’s instructions. The library was sequenced on HiSeq systems (Illumina) using 150 nucleotides (nts) kits and paired-end sequencing.

### SPLiT-Seq data processing

According to the results of FastQC, adaptors or low-quality nucleotides were trimmed by Trim Galore (v0.5.2) using the default parameters. For each paired-end sequencing read, a 10-bp UMI sequence and a 24-bp cell barcode were extracted from the Read 2 file by the tool “preprocess_splitseq.pl” of zUMIs (v0.0.6). The Read 1 was split by different cell barcodes in Read 2 and mapped to the modified *C. elegans* genome (WS263 with an artificial chromosome containing the *C. elegans* version of the GFP sequence) by zUMIs (v0.0.6) and STAR (v2.6.0c). Unique mapping reads were kept. The duplicated reads from the same transcript were excluded based on the UMI information in Read 2. Low-quality transcriptomes were removed from the analysis, yielding 442 single-cell transcriptomes, including transcript counts from 8629 genes. Among these transcriptomes, 41 transcriptomes were assigned to the putative apoptotic cell based on the expression of *egl-1* or *gfp*. Among the total 8624 genes detected in SPLiT-seq, transcripts from 6487 genes were not detected in the putative apoptotic cell population. Enriched GO terms of those 6487 genes were identified using Gene Ontology knowledgebase (Ashburner et al., 2000; Gene Ontology, 2021) (http://geneontology.org/).

### Cell lineage tracing and quantification of reporter expression

Embryo mounting, live imaging, lineage tracing, and expression quantification were performed according to a previously described procedure with minor modifications (Bao and Murray, 2011; Du et al., 2014; Murray et al., 2008). Two- to four-cell stage embryos were collected from young adult worms and mounted between two coverslips in the egg buffer containing 20-30 20-μm polystyrene microspheres and sealed with melted Vaseline (Bao and Murray, 2011). 3D time-lapse imaging was performed at 20 °C ambient temperature using a spinning disk confocal microscope (Revolution XD) for 300-time points at a 75-second interval. For each time point, embryos were scanned for 30 Z focal planes with 1 µm spacing. 3D tiff stack images were processed with the StarryNite software for automated cell identification and tracing to reconstruct embryonic cell lineages using the ubiquitously–expressed mCherry fluorescence. The raw results of cell identification and tracing were subjected to extensive manual inspection and editing using the AceTree program to ensure high accuracy (Du et al., 2014). For each mentioned strain, the cell lineage was traced from the two- or four-cell stage to the 350-cell stage for three or four embryos (experimental replicates). Regions of the chromatids or nuclei are determined by the H2B::mCherry signal. The fluorescent intensity of HDA-1::GFP or LIN-53::mNeonGreen in each traced nucleus at each time point was measured as the average fluorescent intensity of all the pixels within each nucleus and then with the average fluorescent intensity of the local background subtracted (Murray et al., 2008).

### RNA sequencing

Synchronized young adult worms were cultured on RNAi plates for 24 hours. Gravid adults were transferred to new RNAi plates to lay eggs for 1 hour. Eggs were harvested 5 hours post-laid and then lysed with TRIzol reagent (Invitrogen). We extracted the total RNA following the manufacturer’s protocol. The Qubit RNA High Sensitivity Assay Kit (Invitrogen) was used to quantify RNA concentration. The Agilent 2100 bioanalyzer system was used for the assessment of RNA quality. We used samples with an RNA integrity number (RIN) above 6.0 for sequencing library construction. We used identical input total RNA (50 ng to 500 ng) between control and samples for library preparation using the KAPA RNA HyperPrep Kit (KAPA Biosystems, Wilmington, MA, USA). Libraries were analyzed by Agilent 2100 bioanalyzer system for quality control. The library samples were sequenced on an Illumina NovaSeq 6000 platform. Approximately 5 GB of raw data were generated from each sample with 150 bp paired-end read lengths.

### RNA-Seq data analysis

FastQC assessed the quality score, adaptor content, and duplication rates of sequencing reads. Trim_galore (v0.6.0) was used to remove the low-quality bases and adaptor sequences with default parameters. After trimming, paired-end reads with at least 20 nucleotides in length were aligned to the *C. elegans* reference genome (WS263) using STAR (2.5.4b) with default parameters. Read counts representing gene expression levels were calculated with HTSeq (version 0.9.1) using default parameters. Uniquely mapped reads were included. Gene names were annotated on the Wormbase website (https://wormbase.org/tools/mine/simplemine.cgi). DESeq2 package in R programming language was used for differential expression analysis. Differentially expressed genes were defined as upregulated genes (log2-transformed fold change greater than 1) or downregulated genes (log2-transformed fold change less than −1) with a false discovery rate (FDR) less than 0.05. We used the ggplot2 package of R to plot figures.

### Reverse transcription and quantitative real-time PCR

Young adult worms were cultured on RNAi plates for 24 hours. We transferred the gravid adults to new RNAi plates to lay eggs for 1 hour. Eggs were harvested 5 hours post-laid and then lysed with TRIzol reagent (Invitrogen). The RNeasy Mini kit (QIAGEN) prepared the total RNA from ∼20 μl egg pellets for each biological replicate. cDNA from the same amount of RNA between RNAi and control samples were synthesized in a 20 μl reaction volume by the PrimeScript™ RT reagent Kit with gDNA Eraser (TAKARA, Code No. RR047A). One μl of cDNA was used as the template in a 10 μl reaction volume of the PowerUp SYBR Green Master Mix (Applied Biosystems) with four technical replicates. Quantitative real-time PCR was performed in two independent experiments with three biological replicates each time using the Applied Biosystems QuantStudio 1 Real-Time PCR System and normalized to the cell division cycle-related GTPase encoding gene *cdc-42*. Data were analyzed using the standard curve method. Primers used in real-time PCR assays were listed in Extended Data Table 3.

### Chromatin immunoprecipitation

For ChIP assays, chromatin immunoprecipitation was performed as previously described(Mukhopadhyay et al., 2008) with modifications. Mixture stage worms on RNAi plates were harvested and washed three times with M9 buffer. 0.4 ml of packed worms were obtained and frozen into small balls with liquid nitrogen for each ChIP replicate. We quickly ground the little worm balls into powders in liquid nitrogen. Worm powder was crosslinked in 4 ml crosslinking buffer (1.1% formaldehyde in PBS with protease and phosphatase inhibitors) with constant rotation for 15 min at room temperature. Fixed worm samples were quenched by incubation with 0.125 M glycine for 5 min and then washed in cold PBS-PIC (PBS buffer with protease inhibitor cocktail) three times. The tissue pellet resuspended in PBS-PIC was homogenized by 2-3 strokes in a Dounce homogenizer. The cell suspension was centrifuged at 2,000 g for 5 min at 4 °C, and we removed the supernatant.

We performed nuclei preparation, chromatin digestion, chromatin immunoprecipitation, and DNA Purification using SimpleChIP® Enzymatic Chromatin IP Kit (Magnetic Beads) (Cell Signaling Technology, #9003). Samples were sonicated using a Qsonica sonicator (Qsonica Q800R) at 70% amplitude for 5 cycles of 30 sec on and 30 sec off with 4 repeats. The chromatin was digested using 6 μl of micrococcal nuclease (MNase) (CST catalog number 10011S) in 300 μl of buffer B (CST catalog number 7007) containing 0.5 mM DTT for 8 min at 37 °C. The quality of chromatin digestion was analyzed by Agilent 2100 bioanalyzer system. Before IP, the chromatin sample concentration was assessed by the BCA Protein Assay Kit (Tiangen, cat no. PA115-02). 1 μg H3K27ac antibody (Abcam ab4729) or 2 μg IgG was used per IP with 0.5-1 mg input chromatin protein.

According to the manufacturer, we used 200 ng purified DNA samples to construct the ChIP–seq library using the VAHTS Universal DNA Library Prep Kit for Illumina V3 (Vazyme ND607). DNA libraries were sequenced on an Illumina NovaSeq 6000 platform, with 150-bp paired-end sequencing. Three biological replicates were collected for two independent ChIP–seq experiments. For ChIP–qPCR analysis of H3K27ac enrichment in the *egl-1* locus, the same volume of cDNA from the IP experiment described above was used for qPCR analysis. Extended Data Table 3 listed the primers.

### ChIP-Seq data processing

150-bp double-end reads were aligned to the *C. elegans* genome version WBcel235 using Bwa-mem2 version 2.1 with default parameters. Only uniquely mapped reads were kept. Samples were normalized by bins per million mapped reads (BPM) and averaged, and then treated samples were subtracted by input samples to calculate enrichment using deepTools2 (Ramirez et al., 2016). Data were visualized using IGV(Robinson et al., 2011).

### Immunofluorescence

We performed immunofluorescence of *C. elegans* eggs using the freeze-cracking method. In brief, eggs were washed with M9 buffer three times and dripped onto poly-L-lysine coated slides. After freezing in liquid nitrogen for 5 minutes, the coverslip was swiftly removed, and embryos were fixed in methanol at −20 °C for 15 min. We washed the slides twice with PBST (PBS+ 0.05% Tween 20). We incubated slides with 1% BSA in PBST for 30 min at room temperature to block the unspecific bindings. The primary and secondary antibodies were diluted in PBST plus 1% BSA by 1:500-1:2,000 for primary antibodies (ab10799 for histone H3 and ab4729 for H3K27ac) and 1:5,000 for the secondary antibodies (Alexa-488 anti-mouse and Alexa546 anti-rabbit from Invitrogen). The primary antibody was incubated overnight at 4 °C in a moisture chamber, and the secondary antibody was incubated for 1 hour at room temperature. After washing with PBST, we applied 1-2 drops of the Fluoroshield Mounting Medium containing DAPI (Abcam, ab104139) and coverslips to the samples.

### Pharmacological and chemical treatments

To block V-ATPase proton pump activity in the L1-L2 larvae, which have cuticle layers that may compromise agent penetration, the Bafilomycin A1 treatment was designed based on the published protocols (Kumsta et al., 2017; Papandreou and Tavernarakis, 2017; Pivtoraiko et al., 2010). Briefly, gravid worms were placed in 1 drop of ddH_2_O containing 3% (v/v) pelleted OP50 and 100 μM bafilomycin A1 (Abcam, dissolved in DMSO) for 16 h at 20 ℃. After incubation, L1 larvae were washed with M9 buffer and recovered for 5 h on NG agar plates before imaging.

To achieve Q cell conditional HDA-1 depletion, the auxin-inducible degradation (AID) was performed as described previously (Zhang et al., 2015). The natural auxin (indole-3-acetic acid, IAA) was purchased from Alfa Aesar (#A10556). A 400 mM solution in ethanol was stored at 4 °C as stock. Before imaging, L1 larvae were transferred to the S basal buffer supplemented with 3% (v/v) pelleted OP50 and 4 mM of auxin for a 1.5 h treatment.

### Coimmunoprecipitation and Western blot

For immunoprecipitation of endogenous V-ATPase A subunit with HDA-1, HDA-1-GFP knock-in worms were lysed in ice-cold lysis buffer [pH 7.4, 150 mM NaCl, 25 mM Tris-HCl, 10% glycerol, 1% NP-40, 2x cocktail of protease inhibitors from Roche (Complete, EDTA free), 40 mM NaF, 5 mM Na_3_VO_4_]. The soluble fractions from worm lysates were immunoprecipitated with anti-GFP GFP-Trap A beads (Chromoteck, GTA20) for 1 hr at 4 °C. Immunoprecipitates were washed three times with lysis buffer containing 1x cocktail of protease inhibitors from Roche (Complete, EDTA free) and were then transferred to new tubes. Beads were washed again with 100 mM PB (pH 6.0, 6.5, 7.0, or 7.5) and boiled in 1X SDS loading buffer under 95 ℃ for 5min. The protein samples were analyzed by Western blotting.

Protein samples were resolved by SDS–PAGE and transferred to PVDF membranes. Each membrane was divided into two parts to be incubated with GFP (Abcam ab290, 1:4,000) or ATP6V1A (Abcam ab199326, 1:2,000) primary antibodies, followed by HRP-conjugated Goat Anti-Rabbit IgG (H&L) (Easybio, 1:5,000) secondary antibodies. Quantification of the endogenous V-ATPase A subunit interacting with HDA-1 was performed using HDA-1::GFP bands as a calibration standard. Quantitative densitometry of chemiluminescent bands was performed using ImageJ software.

## Data availability

The data sets generated and analyzed in this study are available in the NCBI Gene Expression Omnibus (GEO, http://www.ncbi.nlm.nih.gov/geo) under accession number GSE167379 with the secure token uxwleayqnjkvnkf.

## Acknowledgments

We thank Drs. W. Zhong, G. Garriga, D. Xue, G. Li, Y. Li and for discussion. This study was supported by the National Key R&D Program of China to W.L. and G.O. (2022YFA1302700, 2017YFA0102900, 2019YFA0508401, 2017YFA0503501), and the National Natural Science Foundation of China to W.L., G.O. and Y.C. (grants 32270773, 32070706, 31991190, 31730052, 31525015, 31861143042, 31561130153, 31671451).

## Author Contributions

W. Li, Y. Chai and G. Ou conceived the project. Z. Xie and Y. Chai performed the experiments, collected and analyzed the data with M. Li, Z. Shen, X. Jiang. Z. Zhu. Z. Zhao, L. Xiao, and Z. Du performed the embryonic cell lineage tracing experiment. W. Li, G. Ou and Z. Xie wrote the manuscript.

---

**Figure S1:**
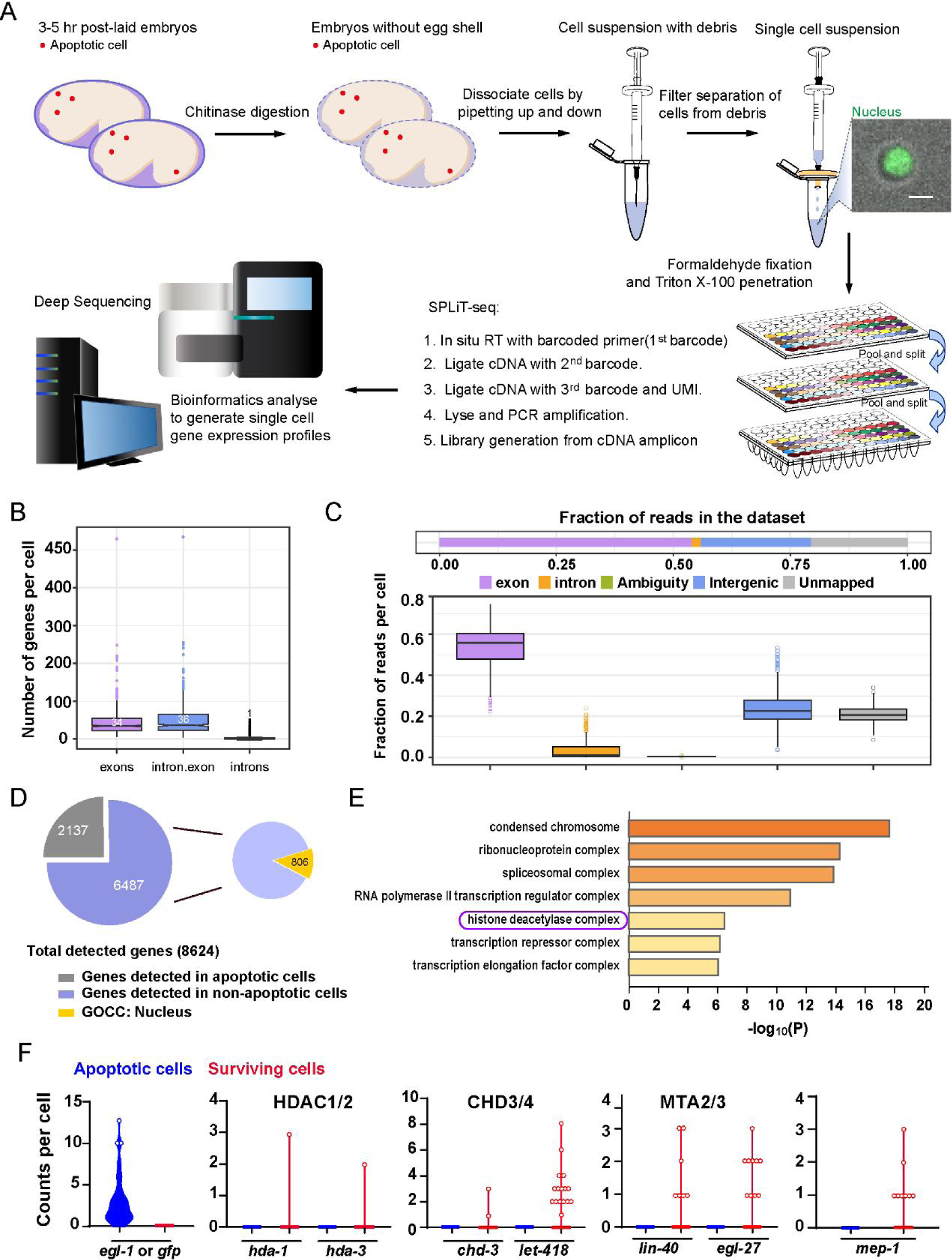
Single-cell sequencing of the *C. elegans* embryonic cells. (A) Schematic of the single-cell SPLiT-Seq workflow for *C. elegans* embryonic cells. (B) Number of genes detected per cell is generated from exon, intron, and exon plus intron mapped reads. Each dot represents a cell, and each box represents the median and first and third quartiles per group. (C) The distribution of reads into different mapping feature categories per cell. Each dot represents a cell, and each box represents the median and first and third quartiles. (D) The left pie chart shows the proportion of genes detected in SPLiT-Seq data from different cell types. The numbers within the pie chart indicate the number of genes. The right pie chart shows that 806 out of 6487 genes, which were not detected in the *egl-1-*positive cells, show enrichment for the nuclear GO terms in the Gene Ontology Cellular Components (GOCC) analysis. See also Supplementary Table 1. (E) Bar plot ranking of representative GOCC terms with a p-value lower than 10^-6^ of 806 genes localized to the nucleus. The “histone deacetylase complex” GOCC term includes *hda-1*, *chd-3*, *lin-40*, and *egl-27*, which encode NuRD subunits. (F) Violin plots show counts per cell for *egl-1*/P*egl-1::gfp* and NuRD component-encoding genes in apoptotic cells and surviving cells.

**Figure S2:**
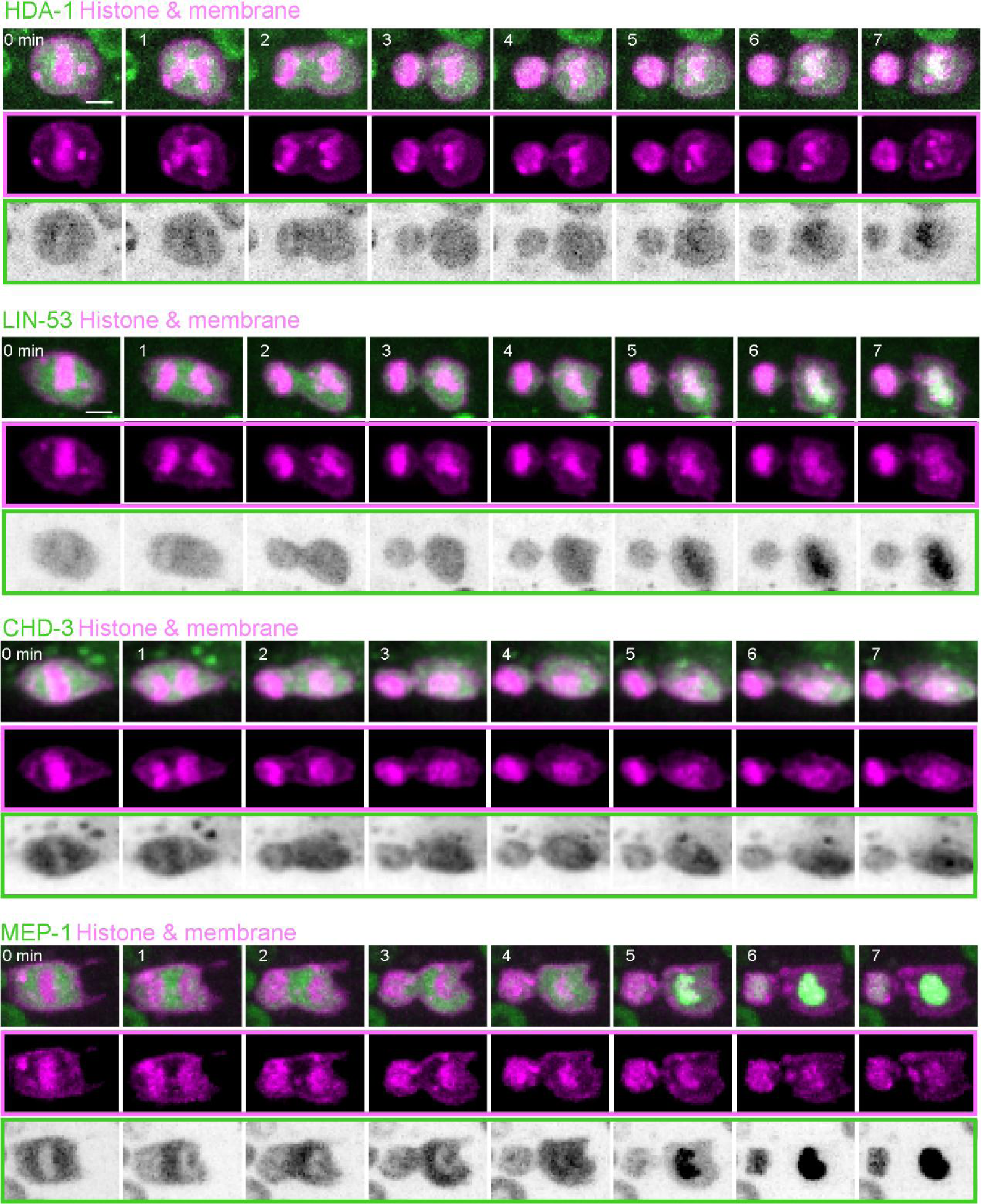
Asymmetric segregation of overexpressed NuRD during ACD of QR.a. Fluorescence time-lapse images of overexpressed GFP-tagged NuRD subunits and mCherry-tagged plasma membrane and histone during asymmetric divisions of QR.a cells. In each panel, the top row shows merged images, the middle row shows mCherry-tagged plasma membrane and histone, and the bottom row shows inverted fluorescence images of GFP. Scale bar: 2 µm. See also in Supplementary Movie S1 and S2.

**Figure S3:**
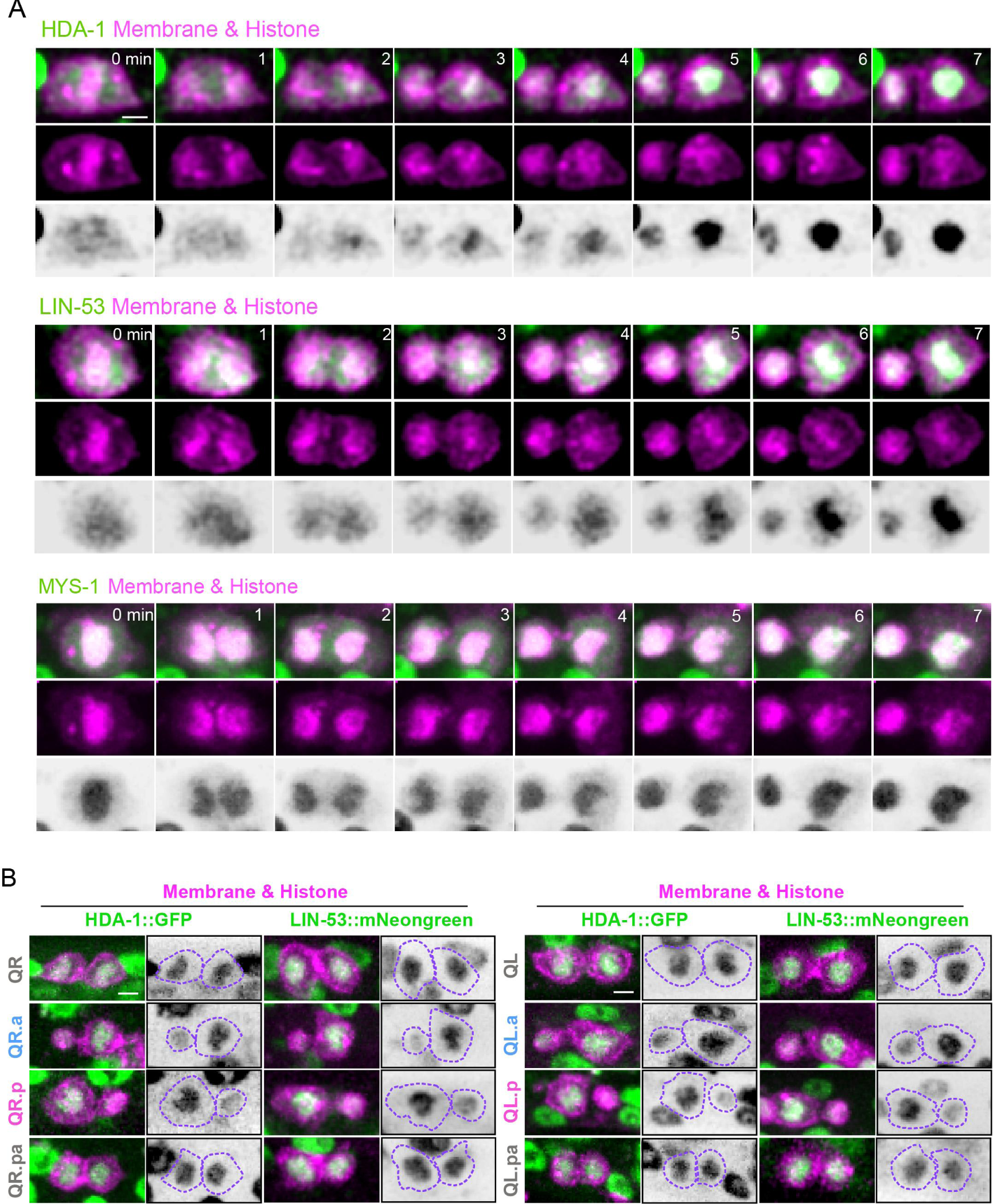
Asymmetric segregation of endogenous NuRD during ACDs of Q cells. (A) Representative images of HDA-1::GFP (top), LIN-53::mNeonGreen (middle), and MYS-1::GFP (down) during QR.a division. The anterior of the cell is on the left. In each panel, the top row shows merged images, the middle row shows mCherry-tagged plasma membrane and histone, and the bottom row shows inverted fluorescence images of GFP/mNeonGreen. Scale bar: 2 µm. See also Supplementary Movie S3-S5. (B) Representative fluorescence images show HDA-1::GFP (left) and LIN-53::mNeonGreen (right) in daughters of indicated Q cells. Inverted fluorescence images of GFP GFP/mNeonGreen signal are on the right of merged images. Dotted purple lines show cell peripheries. The anterior of the cell is toward the left. Scale bar: 2 µm.

**Figure S4:**
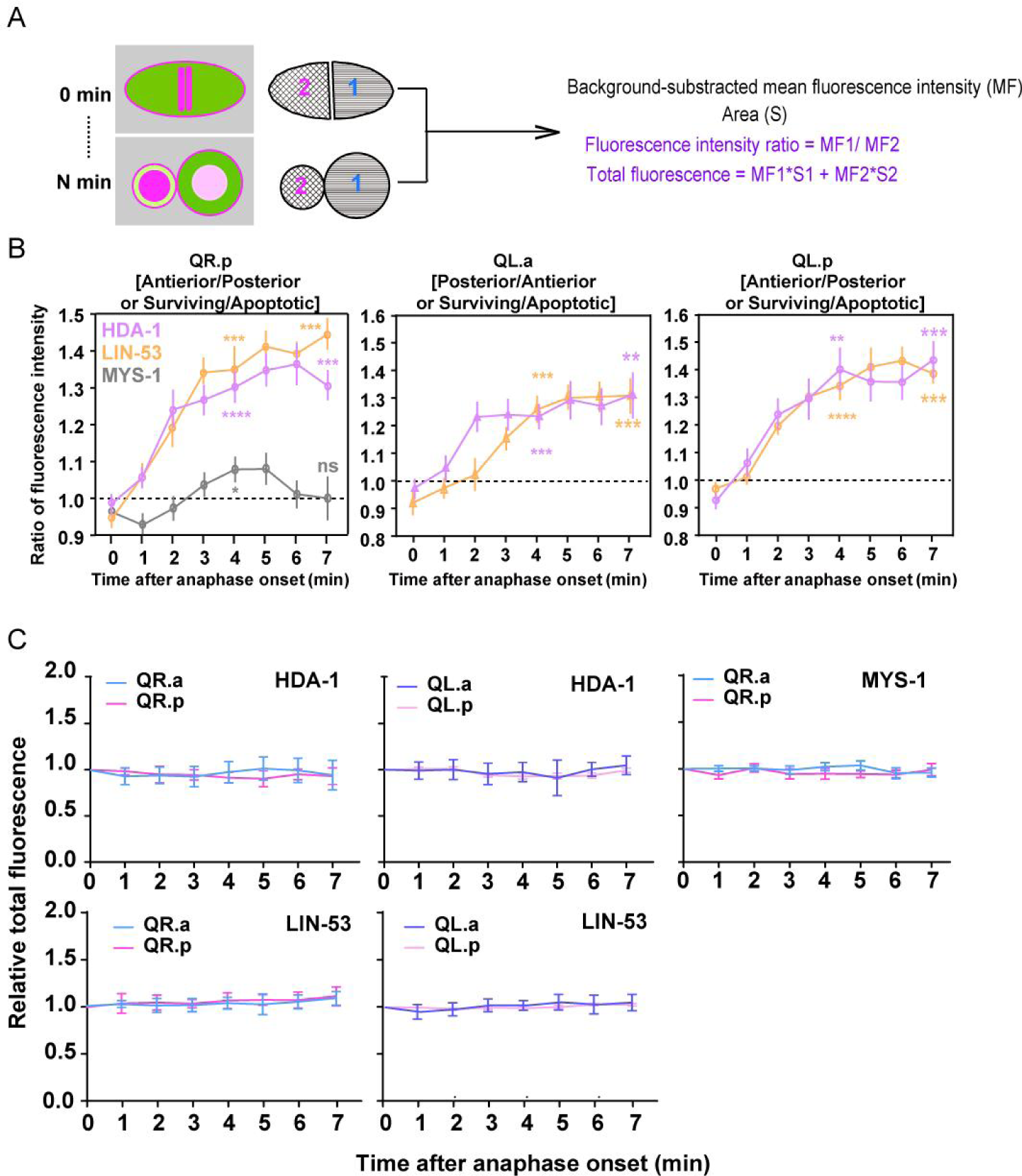
Quantifications of asymmetric NuRD segregation during ACD. (A) Schematics of the fluorescence quantification method. (B) Quantification of the fluorescence intensity ratio (MF1/MF2) changes of HDA-1, LIN-53, and MYS-1 in dividing Q cells. Data are presented as mean ± SEM. N = 10–12. Statistical significance is determined by a one-sample t-test with 1 as the theoretical mean. *p < 0.05, ** p < 0.01, *** p < 0.001, *** p < 0.0001, ns: not significant. (C) The relative total fluorescence of HDA-1, LIN-53 and MYS-1 during ACDs in Figure 1F and S4B. The total fluorescence (MF1*S1+ MF2*S2) at each point was normalized to that at time point 0 min. Data are presented as mean ± SEM. N = 10–12.

**Figure S5:**
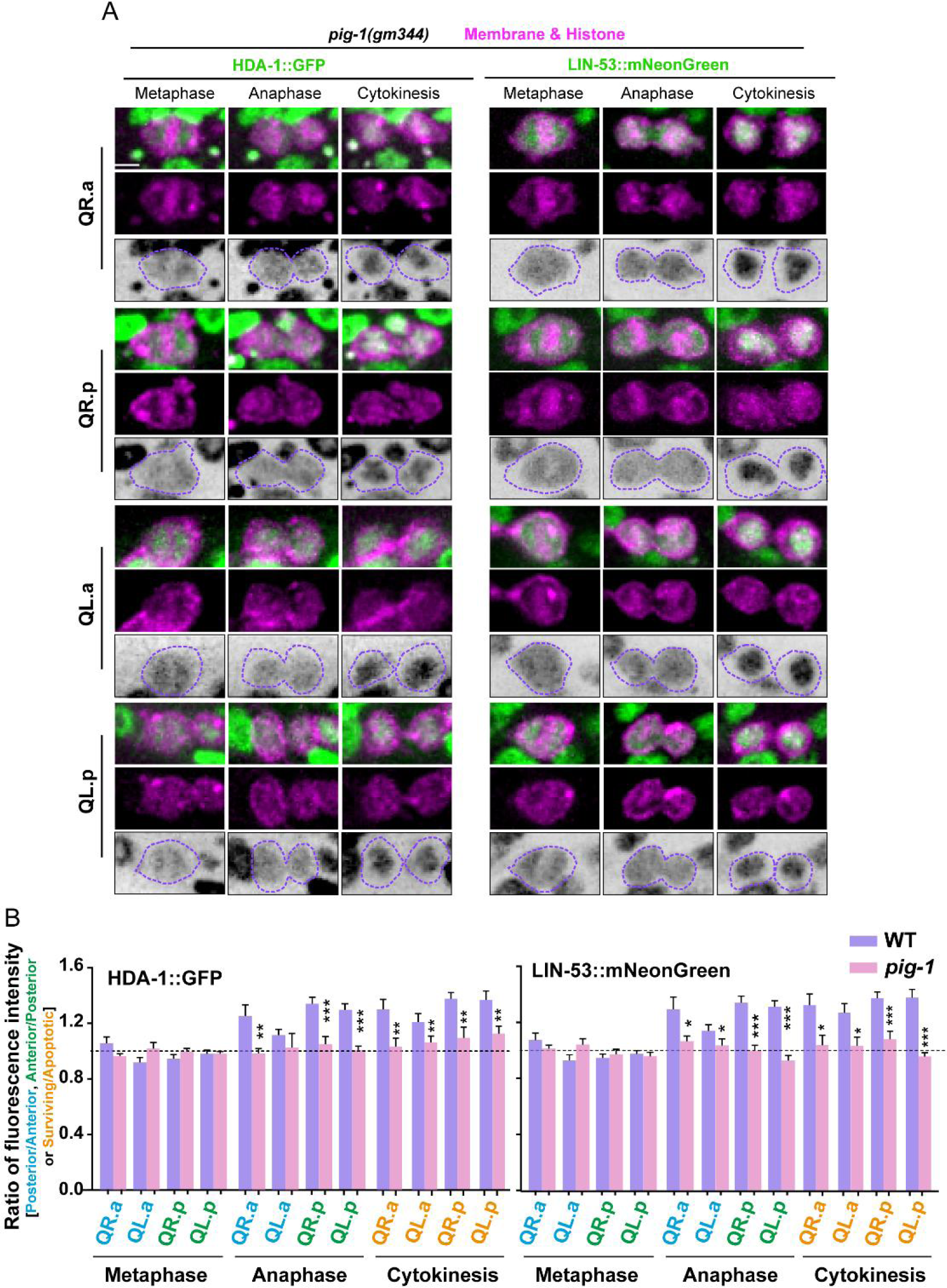
Symmetric NuRD segregation in pig-1 mutant. (A) Representative images of HDA-1::GFP (left) and LIN-53::mNeonGreen (right) during ACDs of Q cells in *pig-1 (gm344)* mutant. In each panel, the top row shows merged images, the middle row shows mCherry-tagged plasma membrane and histone, and the bottom row shows inverted fluorescence images of GFP/mNeonGreen. Anterior of the cell is left. Scale bar: 2 µm. See also Supplementary Movie S6 and S7. (B) Fluorescence intensity ratios of HDA-1::GFP (left) and LIN-53::mNeonGreen (right) between the surviving and apoptotic daughters of Q cells in WT or *pig-1 (gm344)* mutants. Data are presented as mean ± SEM. N = 9–19. Statistical significance is determined by Student’s *t* test. *p< 0.05, **p< 0.01, ***p< 0.001.

**Figure S6:**
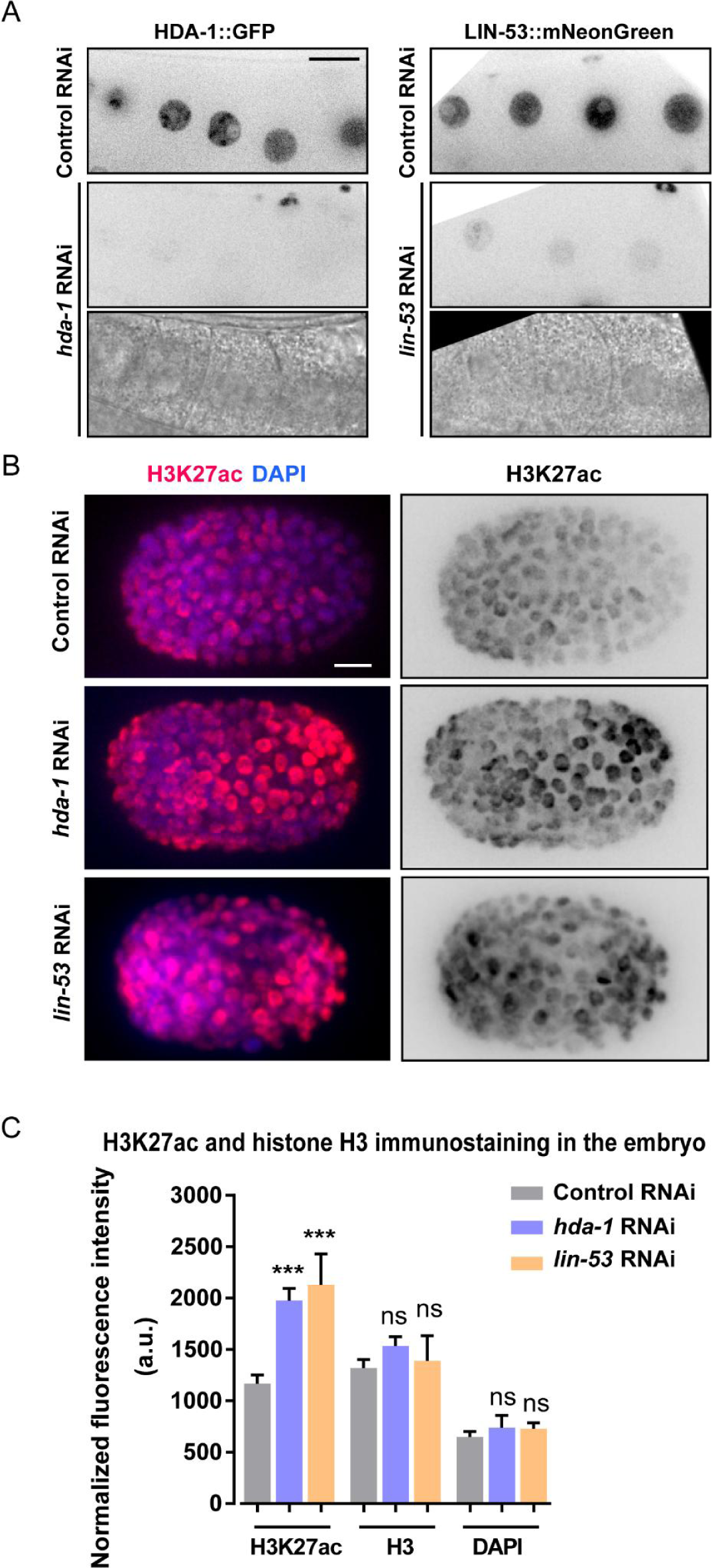
RNAi of *hda-1* and *lin-53* reduce fluorescence of HDA-1::GFP and LIN-53::mNeonGreen and enhance H3K27 acetylation level. (A) Representative inverted fluorescence images (top and middle) and bright-field images (bottom) of oocytes in HDA-1::GFP (left) and LIN-53::mNeonGreen (right) KI animals treated with control, *hda-1* or *lin-53* RNAi. Scale bar: 5 µm. (B) Immunofluorescent images with the anti-H3K27ac antibody of control, *hda-1* or *lin-53* RNAi embryos around the same developmental stage. DAPI stained nuclei. Anterior of the cell is left. Scale bar: 5 µm. (C) Fluorescence intensity quantification of H3K27ac, histone H3, and DAPI from control, *hda-1* or *lin-53* RNAi embryos. The fluorescence intensity is arbitrary units (A.U.). Data are presented as mean ± SEM. N = 16–17. Statistical significance is determined by Student’s *t* test. ***p < 0.001, ns: not significant.

**Figure S7:**
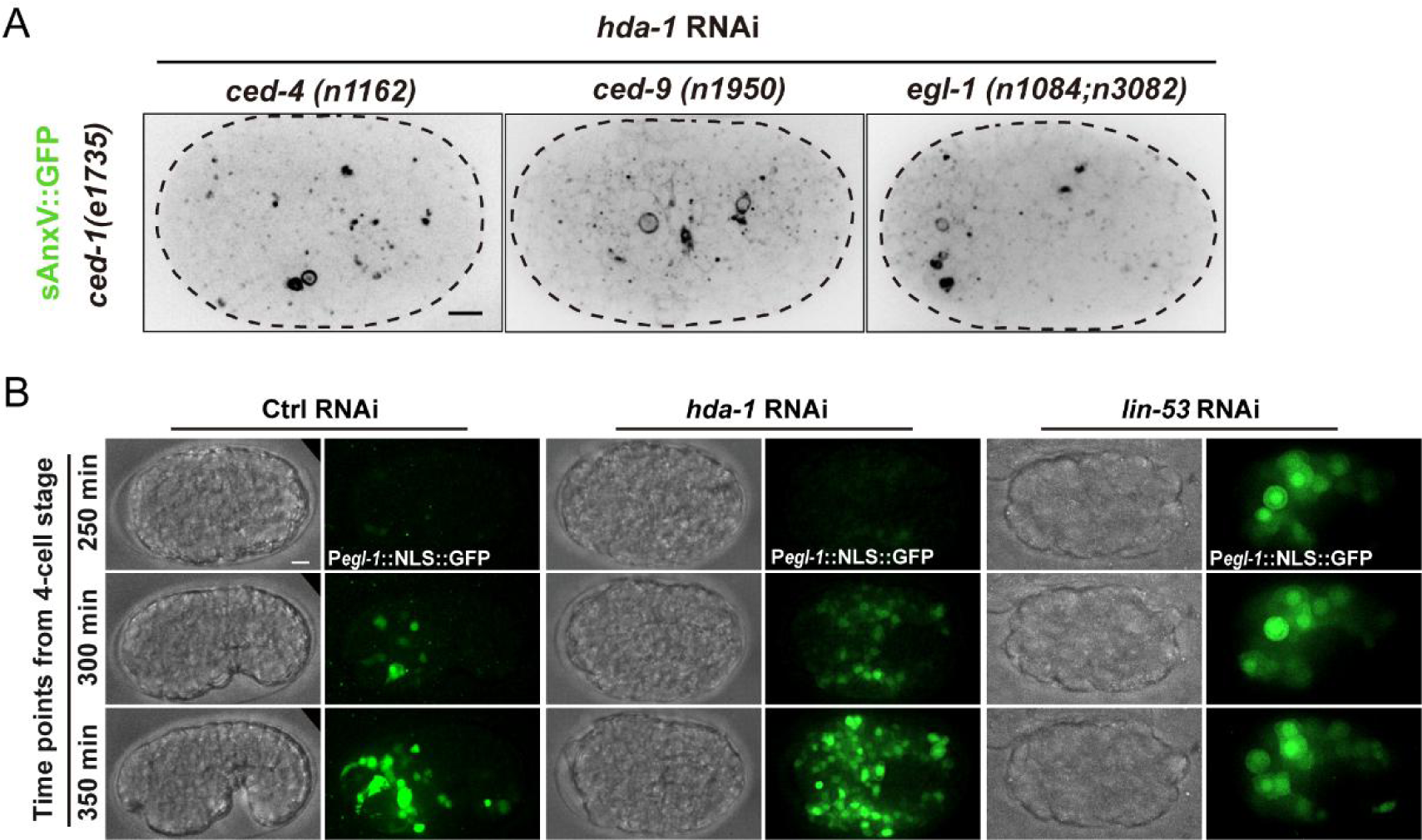
HDA-1 regulates apoptotic cell fate through the canonical apoptosis pathway. (A) Inverted fluorescence images of P*hsp*::sAnxV::GFP in *ced-1(e1735)*, *ced-1(e1735)*; *ced-4(n1162)*, *ced-1(e1735)*; *ced-9(n1950)*, or *ced-1(e1735)*; *egl-1(n1084n3082)* embryos between late gastrulation stage and bean stage, treated with *hda-1* RNAi. Scale bars: 5 μm. (B) Bright-field and fluorescent images of the *hda-1* or *lin-53* RNAi embryos that expressed the P*egl-1*::NLS::GFP reporter. The anterior is to the left, and the dorsal is up. Scale bar: 5 μm.

**Figure S8:**
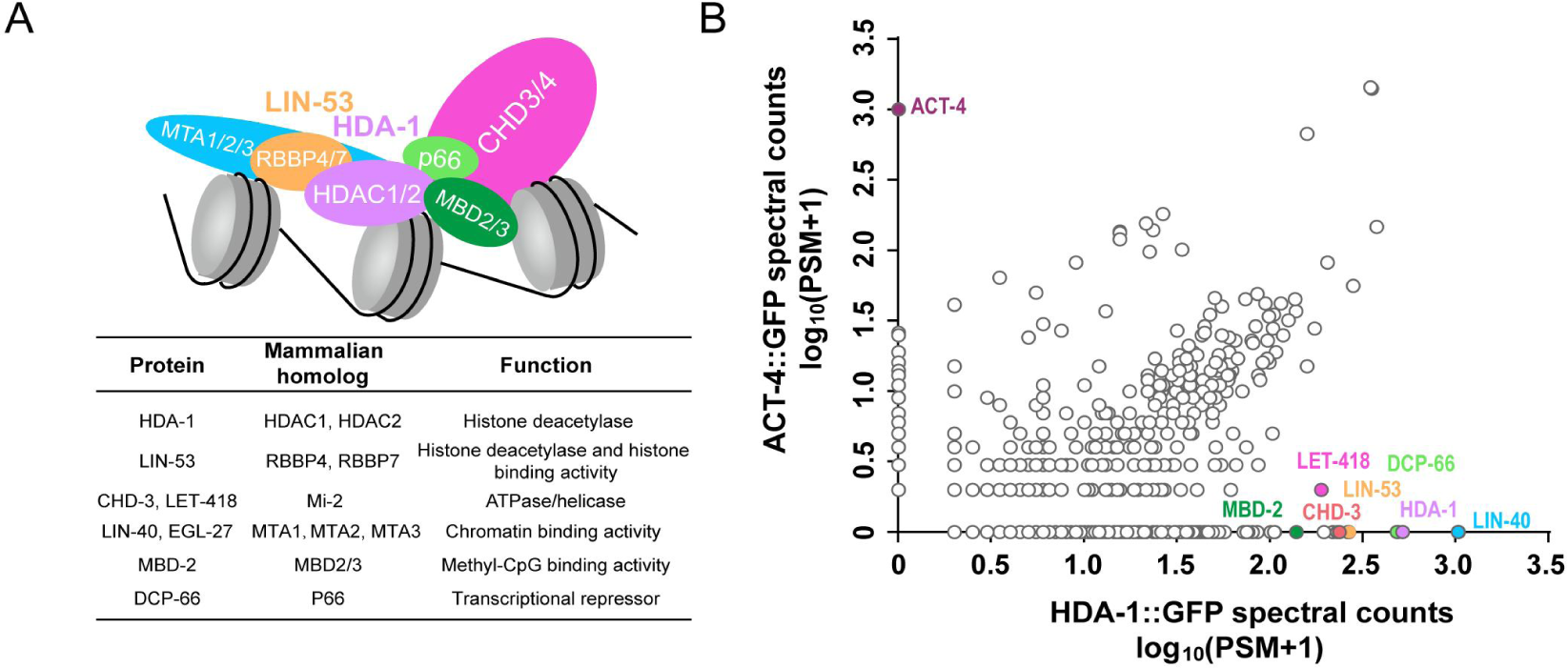
The composition of *C. elegans* NuRD and NuRD subunits identified by co-IP and mass spectrometry. (A) Upper, a schematic representation of the NuRD complex (Bracken et al., 2019; Lai and Wade, 2011). Lower, *C. elegans* homologs of NuRD subunits and their function. (B) Mass spectrometric analysis of proteins purified by anti-GFP agarose beads from HDA-1::GFP KI worms or transgenic worms expressing an actin protein ACT-4::GFP. The plot compares average counts of proteins from two biological replicates co-precipitated with HDA-1::GFP with protein counts co-precipitated with the control ACT-4::GFP. ACT-4::GFP affinity purification pulled down ACT-4 but none of the NuRD subunits (Y-axis), whereas HDA-1::GFP pulled down all the known NuRD components but not ACT-4 (X-axis). The number of PSM is the total number of identified peptide spectra matched for the protein. See also Supplementary Table 4.

**Figure S9:**
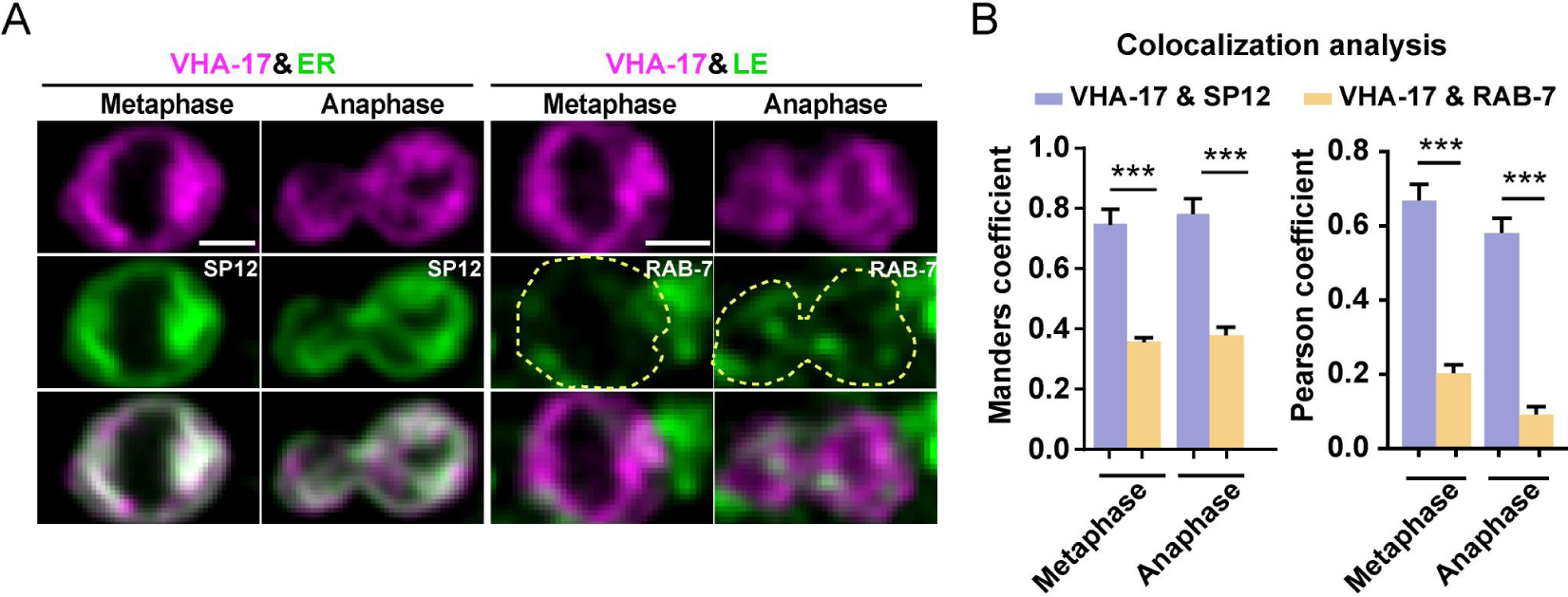
V-ATPase co-localizes with endoplasmic reticulum. (A) Representative double-labeling images of VHA-17 and the ER marker SP12 (left) or the late endosomal marker RAB-7 (right) at metaphase and anaphase in dividing QR.a cells. Scale bar: 2 µm. (B) Quantification of VHA-17 with SP12 or RAB-7 colocalization using Manders overlap coefficient. Data are presented as mean ± SEM. N = 9–13. Statistical significance is determined by Student’s *t* test. ***p < 0.001.

**Figure S10:**
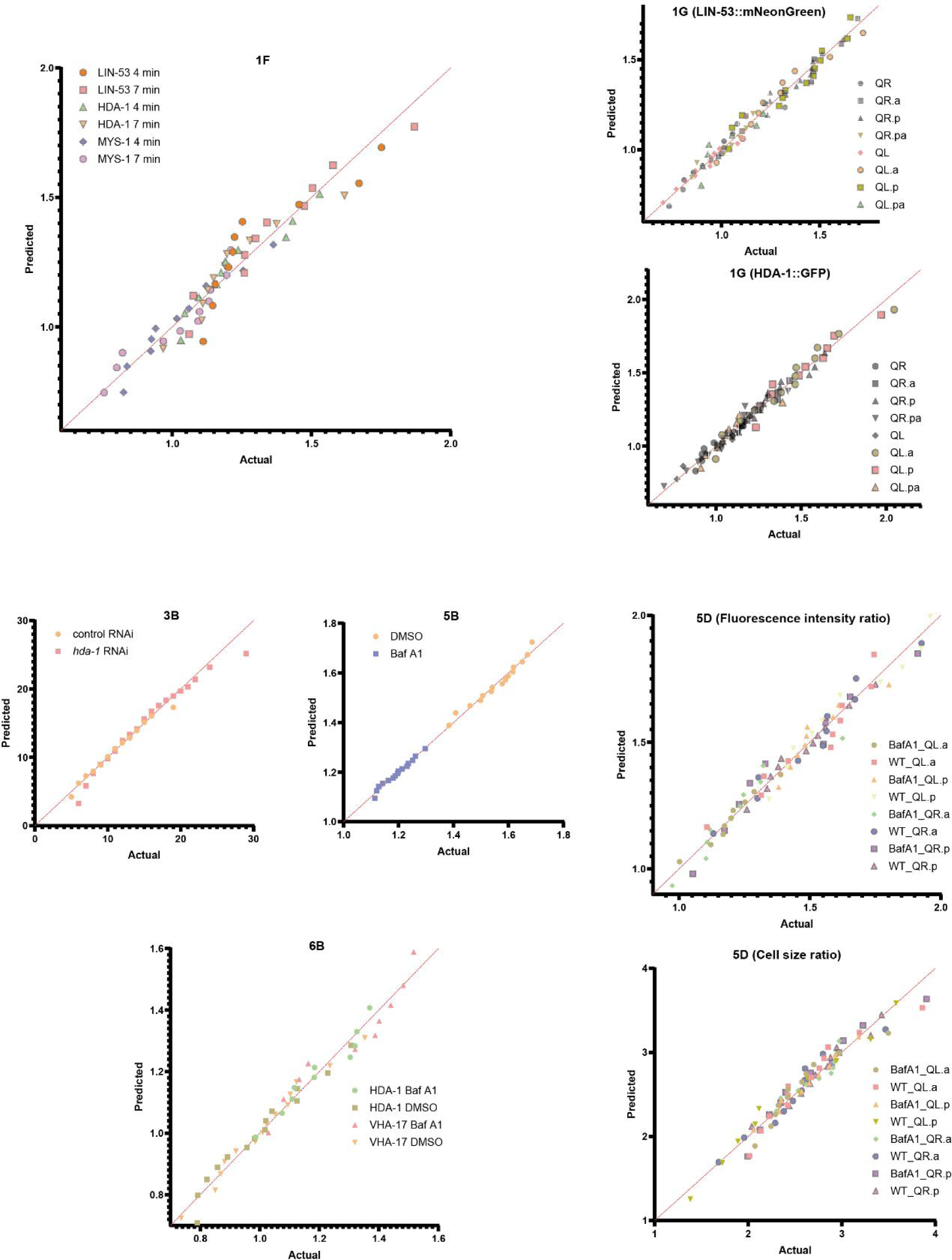
Quantile-quantile (Q-Q) plots for the data in figures 1F, 1G, 3B, 5B, 5D and 6B. The D’Agostino & Pearson and Shapiro-Wilk tests were performed to test the normal distribution of the datasets at 4 and 7 minutes, in figure 1F. The Shapiro-Wilk tests were performed to test the normal distribution of the data in figures 1G, 3B, 5B, 5D and 6B.

**Extended Data Table 1.** *C. elegans* strains in this study.

| Strain Name | Genotype | Method |
| --- | --- | --- |
| N2 | wild-type | N.A. |
| GOU4633 | *cas1133[hda-1::TEV-S::gfp knock-in] V; ujIs113[Ppie-1::H2B::mCherry, Pnhr-2::HIS-24::mCherry, unc-119(+)] II.* | microinjection |
| SYS1031 | *sys1031[lin-53::mNeonGreen knock-in] I; ujIs113[Ppie-1::H2B::mCherry, Pnhr-2::HIS-24::mCherry, unc-119(+)] II.* | microinjection |
| GOU4279 | *cas1133; casIs165[Pegl-17:: myri-mCherry, Pegl-17::mCherry-TEV-S::his-24, unc-76(+)] II.* | Genetic cross |
| GOU4277 | *sys1031; casIs165.* | Genetic cross |
| GOU4636 | *casEX873[Phda-1::hda-1::gfp::unc-54 3'UTR, Pegl-17:: myri-mCherry, Pegl-17::mCherry-TEV-S::his-24].* | microinjection |
| GOU4635 | *casEX874[Plin-53::lin-53::gfp::unc-54 3'UTR, Pegl-17:: myri-mCherry, Pegl-17::mCherry-TEV-S::his-24].* | microinjection |
| GOU4637 | *casEX877[Pchd-3::chd-3::gfp::UNC-54 3'UTR, Pegl-17:: myri-mCherry, Pegl-17::mCherry-TEV-S::his-24].* | microinjection |
| GOU3631 | *wgIs70[mep-1::TY1::EGFP::3xFLAG, unc-119(+)] III; casIs165[Pegl-17:: myri-mCherry, Pegl-17::mCherry-TEV-S::his-24, unc-76(+)] II.* | Genetic cross |
| GOU4634 | *casEX890[Pmys-1::mys-1::gfp-unc-54UTR, Pegl-17:: myri-mCherry, Pegl-17::mCherry-TEV-S::his-24].* | microinjection |
| CU3509 | *ced-1(e1735) I; smIs76[Phsp-16.41::sAnxV::gfp].* | Genetic cross |
| GOU3922 | *ced-3(n2433) IV; ced-1(e1735) I; smIs76[Phsp-16.41::sAnxV::gfp].* | Genetic cross |
| GOU3923 | *ced-4(n1162) III; ced-1(e1735) I; smIs76.* | Genetic cross |
| GOU3924 | *ced-9(n1950) III; ced-1(e1735) I; smIs76.* | Genetic cross |
| GOU3925 | *egl-1(n1084n3082) V; ced-1(e1735) I; smIs76[Phsp-16.41::sAnxV::gfp].* | Genetic cross |
| *smIs89* | *smIs89[Pegl-1::NLS::GFP].* | microinjection |
| GOU4285 | *cas1133; casIs165; pig-1(gm344).* | Genetic cross |
| GOU4281 | *sys1031; casIs165; pig-1(gm344).* | Genetic cross |
| GOU4287 | *cas1133; casEx5309[Phsp-16.2::egl-20, Pegl-17:: myri-mCherry, Pegl-17::mCherry-TEV-S::his-24].* | Genetic cross |
| GOU4283 | *sys1031; casEx5309[Phsp-16.2::egl-20, Pegl-17:: myri-mCherry, Pegl-17::mCherry-TEV-S::his-24].* | Genetic cross |
| GOU4638 | *cas1133; him-5(e1490) V.* | Genetic cross |
| GOU4204 | *sys1031; him-5(e1490) V.* | Genetic cross |
| GOU4607 | *cas1589 [hda-1::GSlinker::degron::TEV-S::gfp knock-in];casIs165; casEx900[Pegl-17:: TIR1::mRuby::unc-54 3'UTR,odr-1::gfp].* | microinjection |
| EX906 | *casEx906 [Pegl-17::vha-17::wormScarlet::unc-54 3'UTR;odr-1::gfp;Pegl-17::TIR1:unc-54 3'UTR] ;cas1589* | microinjection |
| EX913 | *casEx913[Pegl-17::sp12::gfp::unc-54 3'UTR;Pegl-17::vha-17::wormScarlet::unc-54 3'UTR; odr-1::gfp]* | microinjection |
| SYB4702 | *syb4702[gfp::rab-7 knock-in]* | microinjection: syb4702 was generated by Suny Biotech (http://www.sunybiotech.com/) using CRISPR-Cas9. |
| EX909 | *casEx909[Pegl-17::vha-17::wormScarlet::unc-54 3'UTR;odr-1::gfp]; syb4702* | microinjection |
| EX910 | *casEx910 [Pegl-17::super-ecliptic PHluorin::unc-54 3'UTR; odr-1::rfp]; casIs165* | microinjection |
| EX911 | *casEx910; syb4796* | Genetic cross |

**Extended Data Table 2.**
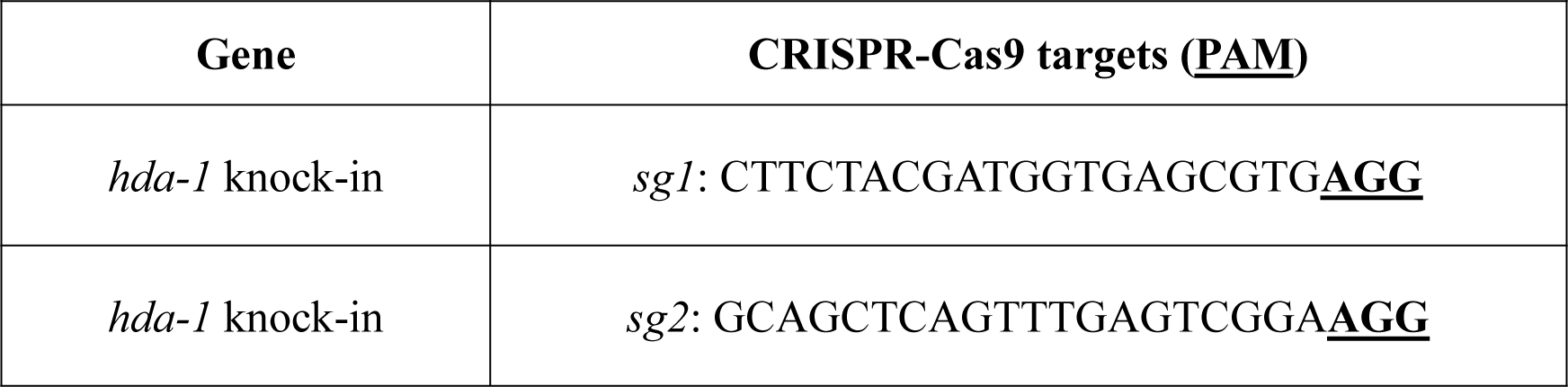
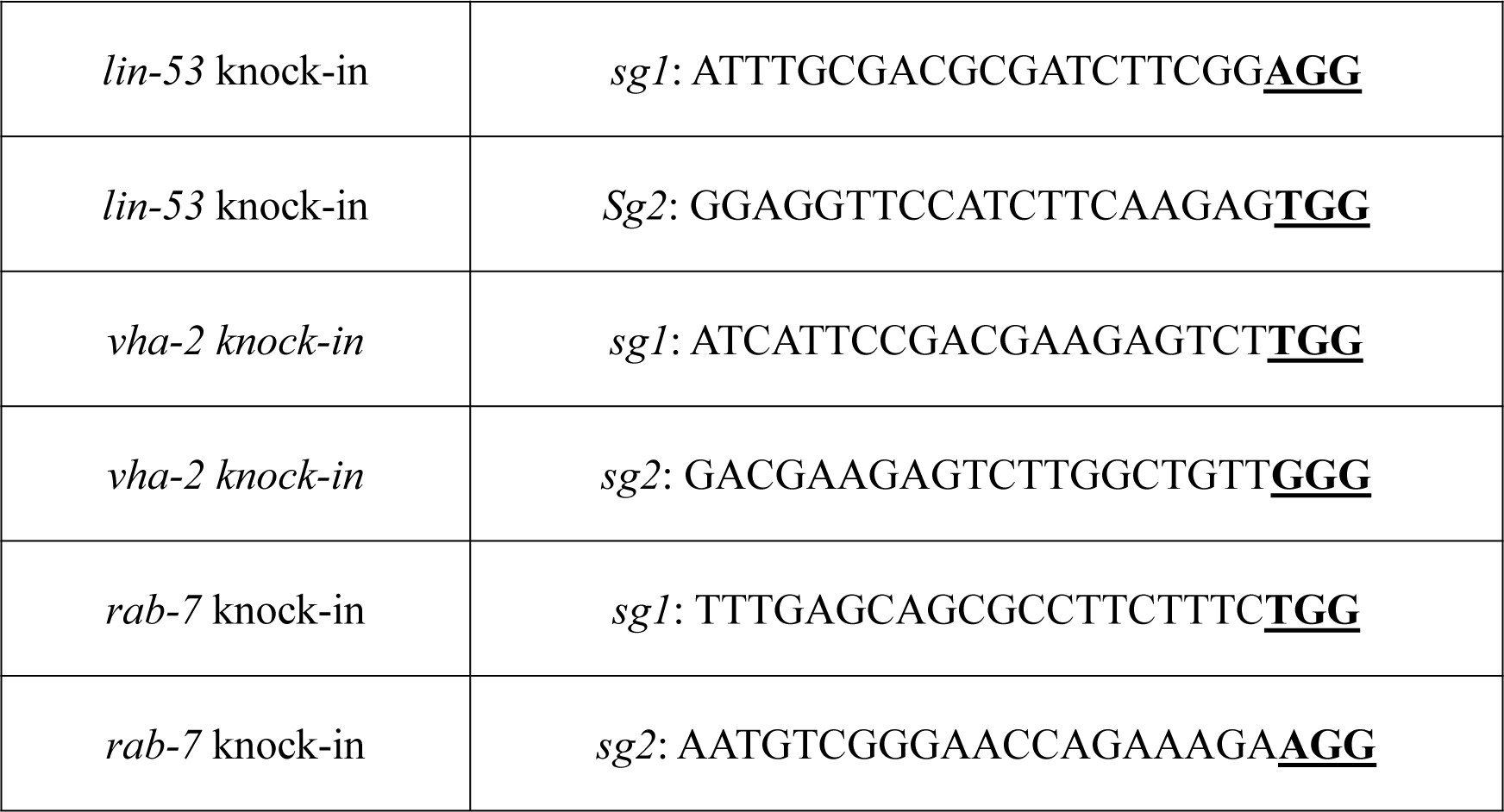
Genomic targets for CRISPR.

**Extended Data Table 3.**
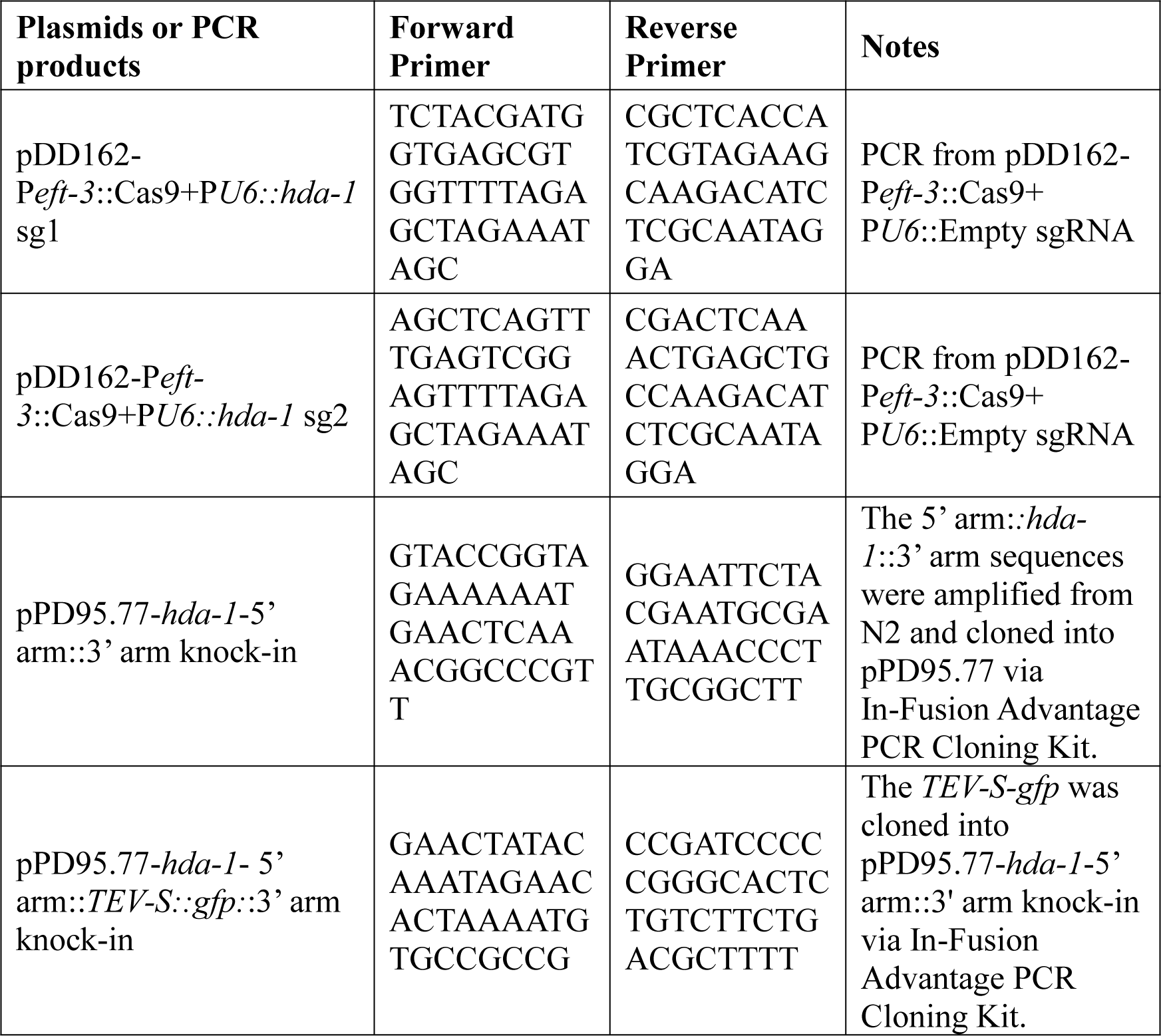
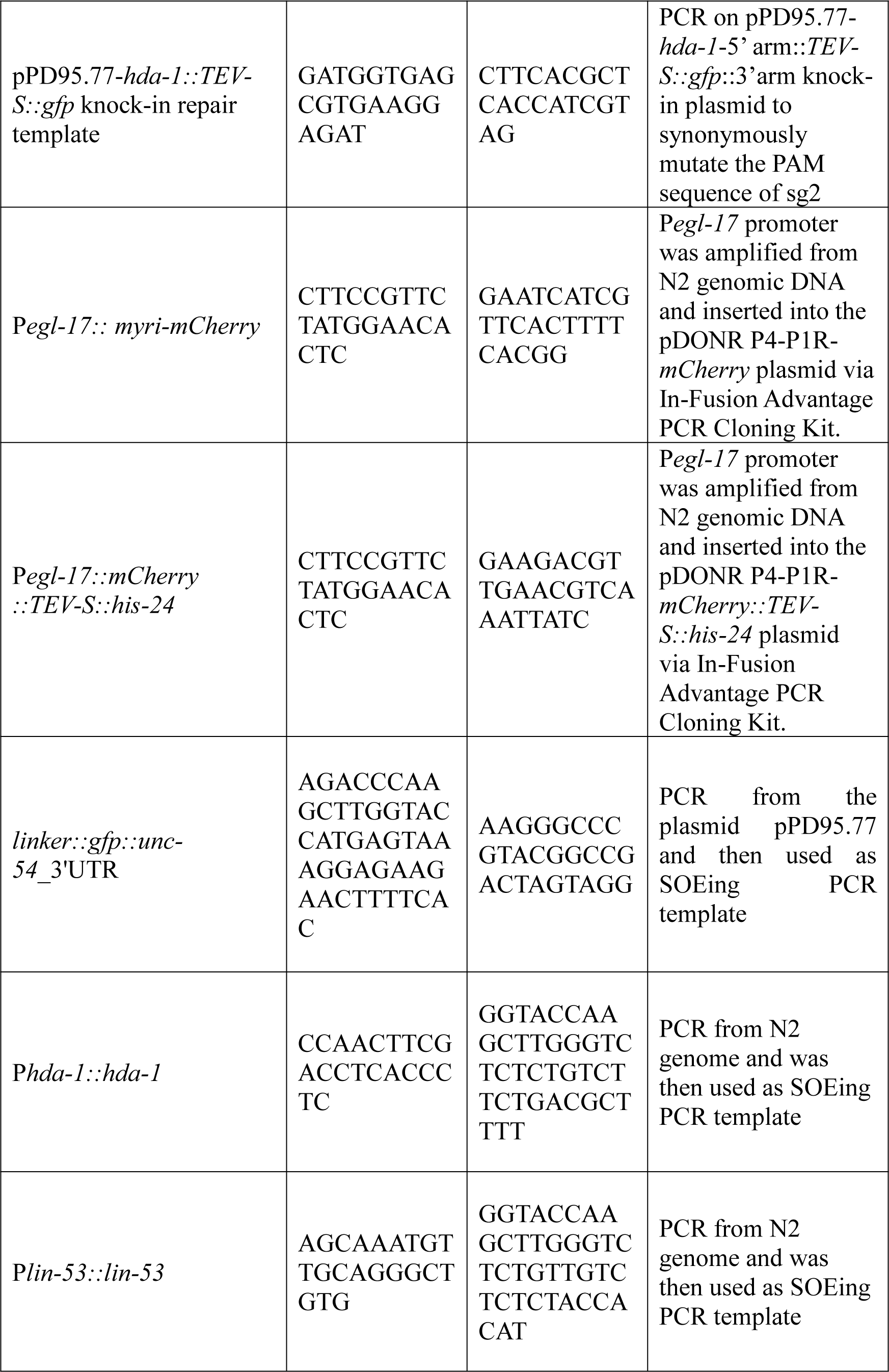
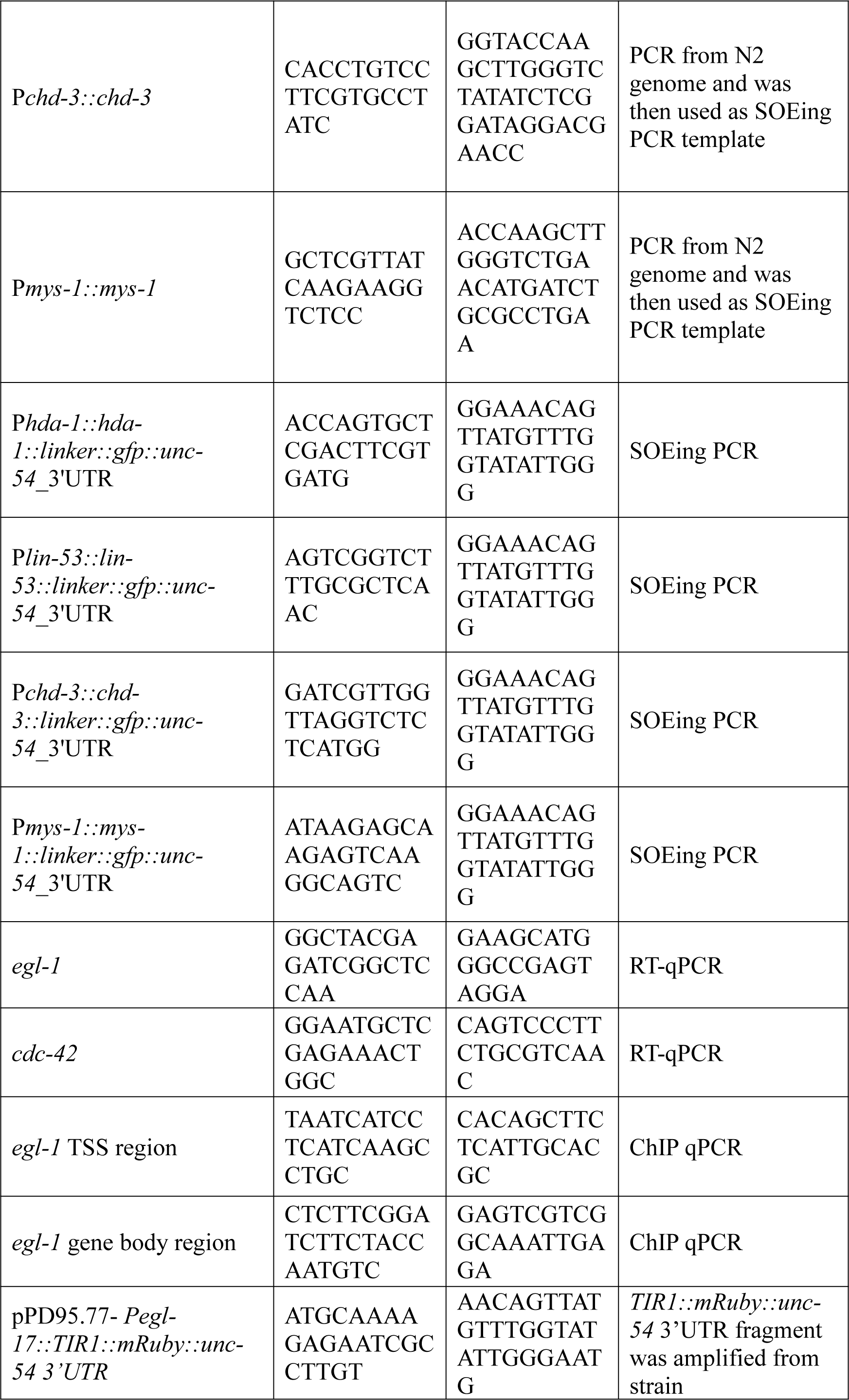
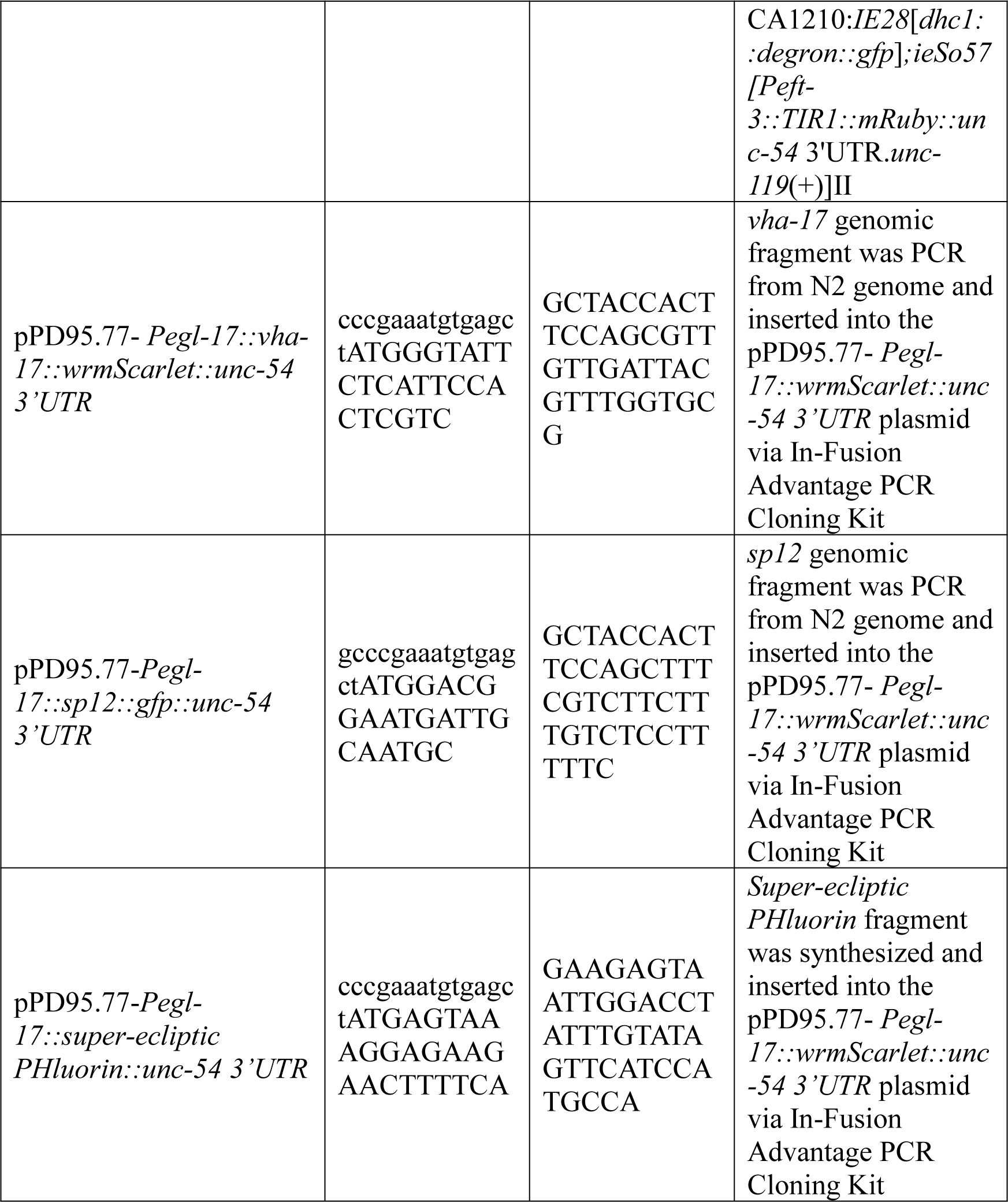
Plasmids and primers used in this study.

## Captions for Supplementary Information

**Supplementary Table 1. Single-cell SPLiT-Seq expression matrix and genes that were not detected from the P*egl-1-NLS-gfp*-positive cells.**

**Supplementary Table 2. The fluorescence intensity ratio of each cell pair and normalized fluorescence intensity of each cell in embryonic lineage tracing assay.**

**Supplementary Table 3. Differential expression of RNA-Seq under control, *hda-1*, or *lin-53* RNAi.**

**Supplementary Table 4. Anti-GFP IP-MS results of HDA-1::GFP KI worms and ACT-4::GFP transgenic worms.**

**Supplementary Movie S1. Dynamics of CHD-3 during QR.a division.** Fluorescence time-lapse movies of CHD-3::GFP (green) and mCherry labeled plasma membrane and histone (magenta) in QR.a. Frames were taken every 1 min. The display rate is 3 frames per second. CHD-3 was asymmetrically segregated into the future surviving QR.ap. Scale bar: 2 μm.

**Supplementary Movie S2. Dynamics of MEP-1 during QR.a division.** Fluorescence time-lapse movies of MEP-1::GFP (green) and mCherry labeled plasma membrane and histone (magenta) in QR.a. Frames were taken every 1 min. The display rate is 3 frames per second. MEP-1 was asymmetrically segregated into the future surviving QR.ap. Scale bar: 2 μm.

**Supplementary Movie S3. Dynamics of HDA-1 during QR.a division.** Fluorescence time-lapse movies of HDA-1::GFP (KI; green) and mCherry labeled plasma membrane and histone (magenta) in QR.a. Frames were taken every 1 min. The display rate is 3 frames per second. HDA-1 was asymmetrically segregated into the future surviving QR.ap. Scale bar: 2 μm.

**Supplementary Movie S4. Dynamics of LIN-53 during QR.a division.** Fluorescence time-lapse movies of LIN-53::mNeonGreen (KI; green) and mCherry labeled plasma membrane and histone (magenta) in QR.a. Frames were taken every 1 min. The display rate is 3 frames per second. LIN-53 was asymmetrically segregated into the future surviving QR.ap. Scale bar: 2 μm.

**Supplementary Movie S5. Dynamics of MYS-1 during QR.a division.** Fluorescence time-lapse movies of MYS-1::GFP (green) and mCherry labeled plasma membrane and histone (magenta) in QR.a. Frames were taken every 1 min. The display rate is 3 frames per second. Anterior, left. MYS-1 was asymmetrically segregated into the future surviving QR.ap. Scale bar: 2 μm.

**Supplementary Movie S6. Dynamics of HDA-1 during QR.a division in the *pig-1* mutant.** Fluorescence time-lapse movies of HDA-1::GFP (KI; green) and mCherry labeled plasma membrane and histone (magenta) during QR.a division in the *pig-1* mutant. Frames were taken every 1 min. The display rate is 3 frames per second. Anterior, left. HDA-1 was evenly segregated into two daughter cells. Scale bar: 2 μm.

**Supplementary Movie S7. Dynamics of LIN-53 during QR.a division in the *pig-1* mutant.** Fluorescence time-lapse movies of LIN-53::mNeonGreen (KI; green) and mCherry labeled plasma membrane and histone (magenta) during QR.a division in the *pig-1* mutant. Frames were taken every 1 min. The display rate is 3 frames per second. Anterior, left. LIN-53 was evenly segregated into two daughter cells. Scale bar, 2 μm.

**Supplementary Movie S8. Dynamics of super-ecliptic PHluorin during QR.a division.** Fluorescence time-lapse movies of super-ecliptic PHluorin (green) and mCherry labeled plasma membrane and histone (magenta) during QR.a division in DMSO and BafA1-treated animals. Frames were taken every 1 min. The display rate is 3 frames per second. Anterior, left. Scale bar, 2 μm.

**Supplementary Movie S9. BafA1 treatment disrupts HDA-1 asymmetry during QR.a division.** Fluorescence time-lapse movies of HDA-1::GFP (KI; green) and mCherry labeled plasma membrane and histone (magenta) during QR.a division in DMSO and BafA1 treated animals. Inverted fluorescence movie of HDA-1::GFP was shown below the merged movie. Frames were taken every 1 min. The display rate is 3 frames per second. Anterior, left. Scale bar: 2 μm.

**Supplementary Movie S10. Dynamics of VHA-17 during QR.a division.** Fluorescence time-lapse movies of wrmScarlet-tagged VHA-17 (magenta) and HDA-1::GFP (KI; green) during QR.a division after DMSO or BafA1 treatments. Inverted fluorescence movie of VHA-17::wrmScarlet was shown below the merged movie. Frames were taken every 1 min. The display rate is 3 frames per second. Anterior, left. Scale bar: 2 μm.

